# DNA walk of specific fused oncogenes exhibit distinct fractal geometric characteristics in nucleotide patterns

**DOI:** 10.1101/2024.07.05.602166

**Authors:** Abhijeet Das, Manas Sehgal, Ashwini Singh, Rishabh Goyal, Mallika Prabhakar, Jeremy Fricke, Isa Mambetsariev, Prakash Kulkarni, Mohit Kumar Jolly, Ravi Salgia

## Abstract

**Background/Objectives:** The complex system of cancer has led to an emphasis on understanding the more general causal relationship within the disease. In this context, concepts of symmetry and symmetry-breaking in distinct biological cell features or components have been examined as an approach to cancer investigation. However, there can be possible limitations in directly interpreting the symmetry-based approach from a physical viewpoint due to the lack of understanding of physical laws governing symmetry in complex systems like cancer.

**Methods:** Fractal geometry and DNA walk representation were employed to investigate the geometric features i.e., self-similarity and heterogeneity in DNA nucleotide coding sequences of wild-type and mutated oncogenes, tumour-suppressor, and other unclassified genes. The mutation-facilitated self-similar and heterogenous features were quantified by the fractal dimension and lacunarity coefficient measures, respectively. Additionally, the geometrical orderedness and disorderedness in the analyzed sequences were interpreted from the combination of the fractal measures.

**Results:** The findings showed distinct fractal geometric features in the case of fusion mutations. It also highlights the possible interpretation of the observed fractal features as geometric analogues concerning explicit observations corresponding to specific cancer types. In addition, the two-dimensional multi-fractal analysis highlighted the presence of a single exponent in the scaling of mutation-mediated gene sequence self-similarity/complexity and heterogeneity.

**Conclusions:** The approach identified mutation-induced geometric features in gene sequences, demonstrating the potential of DNA walks and fractal analysis in translational research regarding cancer. The findings suggest that investigating fractal parameters can capture unique geometric features in nucleotide sequences, contributing to the understanding of cancer’s molecular complexity.

## 1. Introduction

Cancer is a complex system due to interactions between practically immeasurable molecular agents within cells, with other cells in the tumour microenvironment, and across the host organ. It also offers flexibility to deviations in pH, temperature, and fluxes like oxygen, nutrients, etc., along with manifesting structure-independent functionalities (1,2). These observations have led to an emphasis on understanding the causal relationships in cancer to discover and develop more effective approaches to treatment (1,3). In this context, the concept of symmetry-breaking in distinct biological features, i.e., combinatorial (genotypic or phenotypic), geometric (cellular constituents), and functional (interaction among cellular components), has been explored as an approach to understanding cancer (1). It arises from the perspective that symmetry-breaking occurs frequently and is a necessary condition for life (4,5). Nonetheless, symmetry breaking is always incomplete since total breaking would imply total disorder. On the other hand, complete order also cannot make life sustainable as the information would be insufficient for complex biological functions necessary for life. Therefore, this implies that life is a result of perfect synchronization between order and disorder (1,6). However, direct investigation of symmetries, their breaking, and types from a physical viewpoint is difficult in complex systems due to a lack of understanding of the physical laws governing them. Nevertheless, qualitative interpretations about the presence or absence can be made by identifying the orderliness or disorderliness in features of interest. In this context, cancer can be understood as a state characterized by unique patterns of order or disorder. These patterns are defined by the combinatorial, geometric, and functional features in cells and their components, extending beyond normal homeostasis. This deviation from normalcy is reflected in the loss of global functions, observable both from a top-down and bottom-up approach, disrupting the meticulous system required to support life.

Until recently, it was generally held that cancer is a genetic disease (7), and mutations in DNA are the central causal event in carcinogenesis. According to this gene-centric view, the ultimate aim of cell division is to copy the genome of the parent cell precisely and divide it between two daughter cells. During this process, atypical errors can manifest as different types of mutations in specific genes, resulting in genomic alterations in daughter cells. Finally, accumulating these alterations in the resulting genome could cause dysregulation of cell division, disparity between cell growth and death or apoptosis, and ultimately, cancer via disruption of normal cell homeostasis. Notably, genomic alterations have been classified both as a hallmark and driving force in tumourigenesis (8,9). A plentiful of genetic mutations that play a prominent role in cancer pathogenesis have been identified from advancement in cancer genetics subsequently, aiding in discovering unique targeted treatments for patients bearing those mutations in specific cancer types like non-small cell lung cancer (NSCLC) (10,11).

Considering cancer and/or its specific features as a complex system with a specific emphasis on geometric features, the investigation and detection of ordered and disordered states will require consideration of two approaches, namely conventional or Euclidean geometry and fractal or non-Euclidean geometry. Pathologists routinely employ Euclidean geometry to observe distortedness or abnormality in cellular and nuclear structures as they are reliable diagnostic criteria in cancer detection and are also closely related to prognosis (12,13). However, the conventional approach cannot capture the intricate geometric details in complexity and heterogeneity, possibly highlighting states with unique order and disorder states, owing to its variation with the scale of observations. The utilization of fractal geometry can overcome this limitation due to the self-similar characteristics of fractal structures or objects. Nonetheless, until recently, fractal geometry has been sparsely utilized in cancer studies; however, they are recently gaining interest, especially as a criterion for diagnosis and prognosis (14–18).

In fractal geometry, the frequency of occurrence or self-similarity is described by power law in systems/structures exhibiting fractal properties. Consequently, large-frequency events possess a greater probability of occurrence as compared to events with frequencies following a normal distribution. In this context, large deviations in specific cancer phenomena or features can be speculated to be related to fractal properties. Notably, in distinction to mathematical systems, the self-similarity in natural physical systems is statistical and known as self-affinity, and the structures are known as self-affine fractals.

Although there is no rigorous definition or reasoning for the existence of fractal properties in DNA from a biological context, their presence can be qualitatively imagined from the following i) dynamics of evolution and/or functional activity inside the constrained domain (e.g., fractal lung, brain, etc.) might be believed to influence DNA’s spatial geometry (19), ii) the space-filling nature of fractals has been manifested in the untangled or unknotted packing of DNA within cells in three dimensions from the fractal globule structure, and iii) the nucleotide sequences in DNA are not random but follows a pattern that can be repetitive and irregular. Nonetheless, at the DNA level, the assumed fractal structure of gene sequences has been examined using chaos game representation and DNA walks (graphical representation of the nucleotide sequences). The generic aim was to identify the typically opaque long-range correlations in gene nucleotide sequences and disruption reflected by mutation-mediated duplications, repeats, and translocations (14). In particular, considering genomic modifications, DNA walks have been utilized as a model for cancer investigation, diagnostics, and drug resistance analysis (10,14,20,21). Indeed, geometric order and disorder in the specific genes could impact the base pairing (local symmetry) in the genome and may produce alterations in the balance between regular and irregular cell divisions (crucial for biological improvements), which, according to Shahriyari and Komarova (22), can trigger differentiation arrest in cancer growth. Moreover, dysregulation in asymmetric cell divisions can promote oncogenesis. Thus, owing to the statistical scale-invariance fractal characteristics of DNA nucleotide sequences, investigating the trend of fractal parameters can capture the unique qualitative presence of novel geometric features and, subsequently, order, disorder, order-disorder states from the scale of a short segment of sequences to the whole sequence and possibly to the complete genome.

Nevertheless, to the best of our knowledge, there are limited, or no report(s) emphasizing geometrical features and states the analysis of DNA nucleotide sequences based on the non-Euclidean approach via combining both mono- and multi-fractal techniques, respectively in cancer research. Motivated by the aforementioned we have utilized image analysis and fractal geometry to investigate the mutations-induced geometric features (order and disorder), highlighting the nucleotide sequence or genomic structural complexity and heterogeneity in wild-type (proto-oncogenes, tumour suppressor genes, and other unclassified genes) and their mutated variants related to diverse classes of cancer using the DNA walk representation. The assumptions of the performed study are as follows: [1] Geometry signifies the spatial arrangement and distribution of nucleotides in the gene sequence, [2] Correlation exists in gene sequences since the DNA chain is not one-dimensional; however, the molecules of DNA are arranged in a complex three-dimensional structure where distance nucleotide base pairs are not only in close geometrical proximity but can also propagate along the chain in addition to jumping several steps along the chain (23). This correlation can be interpreted in the context of fractal dimension (FD) since it is correlated to the Hurst Exponent (H = 2 – FD), via the box-counting algorithm) (24), [3] nucleotide sequences evolve in far-from-equilibrium conditions along with natural selection, genetic drift, etc. thus, are not random structures, and [4] the non-white pixel distribution in the images represents the distribution of nucleotides in the investigated genes.

## 2. Materials and Methods

The CoDing Sequence (CDS) for wild-type genes was downloaded in FASTA or text file format from the NCBI RefSeq Database. The studied list of genes, considered chromosome locations, and RefSeq ID are listed in Table 1. The CDS Sequences for fusion genes were obtained from the FusionGDB database. The point mutations and deletion (DEL)-insertion (INS)-deletion/insertion (DELINS) mutations were created by manually changing the codons at specified locations for common mutations reported in pieces of literature.

**Table 1.** Chromosomal location and the RefSeq ID for the wild-type variant of the studied oncogenes.

| Gene | Chromosome Location | RefSeq ID |
| --- | --- | --- |
| EML4 | 2p21 | NM_019063.5 |
| EGFR | 7p11.2 | NM_005228 |
| ALK | 2p23.2-p23.1 | NM_004304.5 |
| BAG4 | 8p11.23 | NM_004874 |
| FGFR1 | 8p11.23 | NM_023110 |
| RAD51 | 15q15.1 | NM_002875.5 |
| MET | 7q31.2 | NM_000245 |
| KIF5B | 10p11.22 | NM_004521 |
| PML | 15q24.1 | NM_033238 |
| RARA | 17q21.2 | NM_000964 |
| ATF1 | 12q13.12 | NM_005171 |
| EWSR1 | 22q12.2 | NM_005243 |
| SFPQ | 1p34.3 | NM_005066 |
| TFE3 | Xp11.23 | NM_006521 |
| TMPRSS2 | 21q22.3 | NM_005656 |
| ETV4 | 17q21.31 | NM_001079675 |
| PTPRK | 6q22.33 | NM_002844 |
| RSPO3 | 6q22.33 | NM_032784 |
| MYB | 6q23.3 | NM_001130173 |
| NFIB | 9p23-p22.3 | NM_001190737 |
| BRAF | 7q34 | NM_004333 |
| KIAA1549 | 7q34 | NM_001164665 |
| CRTC1 | 19p13.11 | NM_015321 |
| MAML2 | 11q21 | NM_032427 |
| FLI1 | 11q24.3 | NM_002017 |
| SS18 | 18q11.2 | NM_001007559 |
| SSX1 | Xp11.23 | NM_005635 |
| SSX2 | Xp11.22 | NM_175698 |
| TP53 | 17p13.1 | NM_000546.6 |
| ESR1 | 6q25.1-q25.2 | NM_000125.4 |
| KRAS | 12p12.1 | NM_004985 |
| PIK3CA | 3q26.32 | NM_006218.4 |

The DNA walks were generated according to the protocol reported by Hewelt *et al.* (10). Briefly, for the obtained DNA gene sequences, the FASTA files were read using the SeqIO of Bio module, and the parse function and walks were generated using the turtle library in Python. The directional encoding was A(west), T(east), C(south), and G(north). In the case of fusion, all the genes were color-coded and SequenceMatcher from the Difflib library to separate the gene components. The as-generated walks were saved as scalable vector graphics (svg) files using the canvasvg library and later converted to joint photographic experts group (jpeg) format for mono- and multi-fractal analysis.

Preceding analysis of as-generated DNA walks, the pixel columns and rows of individual images (for each group of wild-type and their mutated variants) were standardized, the contrast was normalized, and possible noise was removed without removing the edges from the images. Subsequently, the mono-fractal measures i.e., FD and Lacunarity Coefficient (LC) of walks corresponding to the wild-type and mutated oncogenes performed using the Fraclac plugin of ImageJ software. The two-dimensional (2D) box-counting algorithm and sliding box-counting algorithm were employed for computing the FD and LC measures, respectively. The 2D multi-fractal analysis was performed from the MultiFrac plugin of ImageJ using the box-counting algorithm for binary and grayscale images (25). The computed values of mono- and multi-fractal measures, and the graphical representation of scaling of mass exponent with moment are given in the Supplementary Material.

## 3. Results

### 3.1 Mono-fractal Analysis

#### 3.1.1 DNA walks of wild-type genes

DNA walks of the genes listed in Table 1 were generated in a 2D space in their wild-type (WT) forms, and the FD and LC values were computed using image analysis. The variation in FD and LC is shown in a scatter plot in Fig. 1 for the investigated WT genes, and their parametric values are recorded in Table S1 (Supplementary Material). Here, we did not perform correlation analysis between the parameters since they are mathematically exclusive and quantify different geometric aspects, i.e., complexity and heterogeneity in a complex system. The studied WTs represented different types of cancer i.e., lung adenocarcinoma (EML4, EGFR, ALK, BAG4, FGFR1, RAD51, MET, KIF5B), breast carcinoma (PML, RARA, ESR1), renal carcinoma (ATF1, EWSR1, SFPQ, TFE3), prostate cancer (TMPRSS2, ETV4), colorectal cancer (PTPRK, RSPO3), synovial sarcoma (SS18, SSX1, SSX2) and possible involvement in other carcinomas and sarcomas (MYB, NFIB, BRAF, KIAA1549, CRTC1, MAML2, FLI1, TP53, KRAS, PIK3CA).

**Fig. 1.**
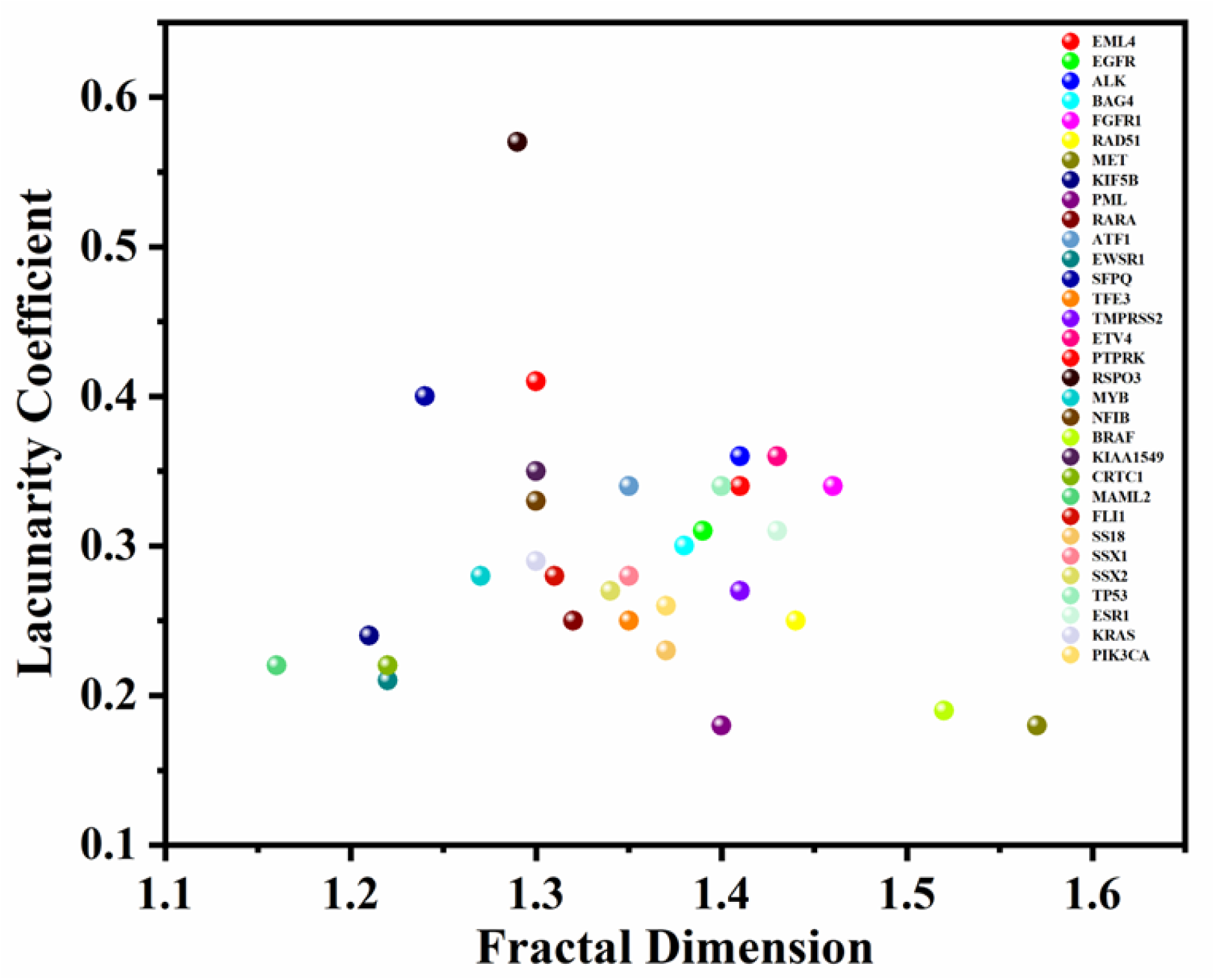
Scatter plot displaying the variation in fractal dimension and lacunarity coefficient for the studied genes in their wild-type variants.

Fractal structures display a repeated pattern at every scale of resolution, i.e., scale invariance where their FD dictates the relationship between the measure and the scale. FD is also used as a measure of complexity where a higher (lower) FD value will imply a greater (lesser) complexity in the investigated system’s feature. However, different spatial patterns have been reported to display a similar value of FD (26). In this regard, another metric, i.e., fractal lacunarity, which measures the distribution and size of voids and/or translational (or rotational) invariance in a texture/pattern or an image and whose strength is determined by the LC can be used along with FD to avoid ambiguity in results. In the context of a DNA gene sequence walk in a 2D space, the value of FD will lie between 1 and 2 where, FD ≈ 1 will represent a non-fractal or conventional one-dimensional Euclidean signature of the sequence whereas, any comprehensible increase from 1 will indicate enhancement in geometrical complexity and self-similarity (order). On the other hand, a larger (smaller) lacunarity coefficient, in the range of 0-1, will imply a lower (higher) rate of alternating base pair patterns consequently, disorderliness (orderliness) in geometry. Finally, in the biological and geometrical context, their combination, i.e., small LC and large FD, will imply low heterogeneity and high self-similarity in nucleotide base pair variations, thus, ordered geometry and vice-versa.

The computed values of FD and LC corresponding to the individual coding sequences of analyzed WT genes are listed in Table S1. It is noteworthy that the FD and LC values of the maximum number of sequences lie in the range of 1.2-1.4 and 0.2-0.4, respectively. However, interesting outliers are observed in the case of MET and BRAF (or B-Raf) proto-oncogenes displaying FD ≈ 1.57 & 1.52 with LC ≈ 0.18 & 0.19, respectively. This suggests an enhanced rate of alternating base pair patterns (14) subsequently, ordered spatial geometry of nucleotide distribution pattern. Nevertheless, the overall trend in variation of FD and LC parameters might imply the existence of a unique type of symmetry, i.e., scaling symmetry in the CDS of WT proto-oncogenes and tumour suppressor genes.

In investigations related to spatial and/or temporal correlation, the Hurst exponent (H), H > 0.5, H < 0.5, and H = 0.5 indicates persistent, anti-persistent, and random behavior, respectively of a spatial pattern (24). Subsequently, for this study, in consideration of its relationship with FD, as mentioned earlier, FD < 1.5, FD > 1.5, and FD = 1.5 will imply a long-range correlation, short-range correlation, and no correlation (white noise), respectively among nucleotides in a DNA walk. Nonetheless, it should be noted that here, long (short)-range correlations do not signify the effect of a segment of the base pair over another segment thousands of base pairs away (memory effect) but imply similar (dissimilar) variation in base pair density in gene sequences at distant lengths (27). In this study, a long-range correlation was realized for most of the sequences while a short-range correlation was measured for MET and BRAF.

#### 3.1.2 DNA walks of mutated oncogenes and tumour-suppressor gene

The walks for mutated (fusion, point mutation, insertion (INS), deletion (DEL), deletion-insertion (DELINS)) oncogenic and tumour-suppressor variants of the respective WT counterparts were generated, and their FD and LC were computed from image analysis. The variation of FD and LC corresponding to the WT and their mutant versions are presented in Fig. 2 and 6. The parametric values are tabulated in Tables S2(a) and S2(b), respectively. In addition, for identification and investigation of the unique ordered disordered geometric states in mutated oncogenes, the complexity/self-similarity and heterogeneity of WT nucleotide sequences were taken as the reference or control.

**Fig. 2.**
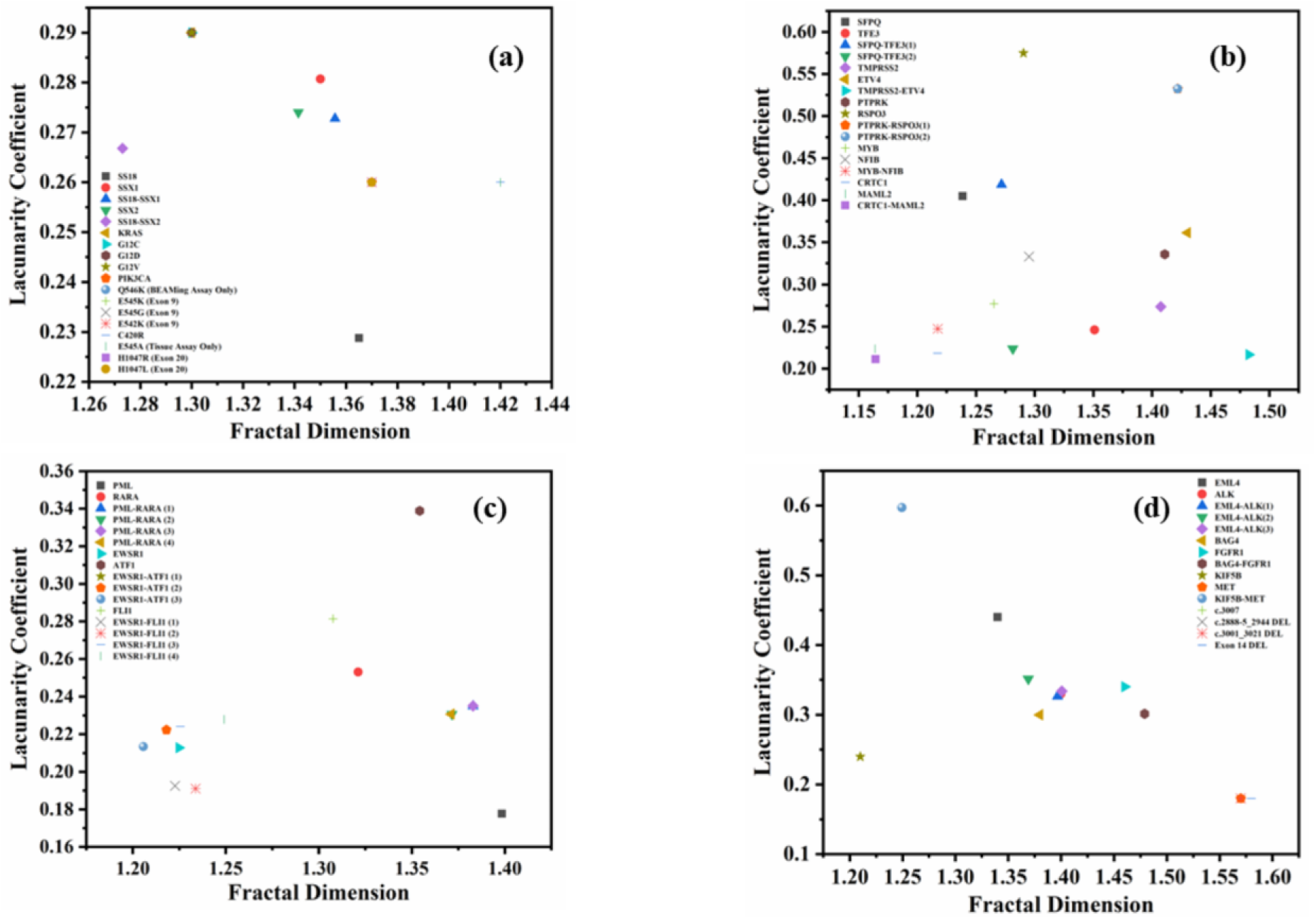
Variation in fractal dimension and lacunarity coefficient in **(a)** wild-type of SS18, SSX1, SSX2, KRAS, and PIK3CA along with their respective fusion and points mutations, **(b)** wild-type of SFPQ, TFE3, TMPRSS2, ETV4, PTPRK, RSPO3, MYB, NFIB, CRTC1, and MAML2 along with their respective fused variants, **(c)** wild-type of PML, RARA, EWSR1, ATF1, and FLI1 with their respective mutated forms, and **(d)** wild-type of EML4, ALK, BAG4, FGFR1, KIF5B, and MET along with their forms with fusion, point-, and deletion mutation.

The variation of mono-fractal measures for SS18, SSX1, and SSX2, and their fused variants (SS18-SSX1) and (SS18-SSX2), are shown in Fig. 2(a). The figure also displays the variation of FD and LC for WT KRAS and PIK3CA along with their point mutated variants. A lower degree of geometric self-similarity and long-range correlation was noted for SS18 and SSX1, individually and subsequently, for SS18-SSX1. A similar case was also observed for SS18 and SS18-SSX2, respectively. The observations suggest no loss in geometric order in WT genes due to fusion. The results were similar for KRAS and its point-mutated variants (G12C, G12D, and G12V). For PIK3CA, low variance in base pairs alterations, resulting from high FD and low LC along with spatial long-range correlation, was observed. In addition, the geometric order of the WT gene was not lost due to the investigated point mutations. Fig. 2(b) displays the variation in FD and LC for wild-type SFPQ, TFE3, TMPRSS2, ETV4, PTPRK, RSPO3, MYB, NFIB, CRTC1, MAML2, correspondingly along with their respective fusion mutated versions. A lower degree of statistical self-similarity and high translational variance, and thus, spatial heterogeneity and, consequently, disorder geometry, was highlighted by SFPQ while the opposite was true for TFE3. Interestingly, their two fused variants, namely, SFPQ-TFE3(1) and SFPQ-TFE3(2), displayed geometric characteristics of SFPQ and TFE3, respectively, suggesting the occurrence of disorder in TFE3 geometry for SFPQ-TFE3(1), and in SFPQ for SFPQ-TFE3(2), respectively. With regard to TMPRSS2, a high FD and low LC were observed indicating geometric order whereas, ETV4 showed augmented geometrical disorder with large FD (high complexity) and LC (high heterogeneity). Interestingly, TMPRSS2-ETV4 revealed considerable decrement in the variance of nucleotide base pair patterns consequently, high geometrical order; however, short-range correlation among base pairs was observed. The respective DNA walks corresponding to WT TMPRSS2 and ETV4 and their fused versions are shown in Fig. 3 with the mutated region color-coded. A high (low) self-affinity and low (high) spatial heterogeneity were observed for PTPRK (RSPO3). However, increasing complexity and heterogeneity in nucleotide base pair patterns were observed in PTPRK-RSPO3(1) and PTPRK-RSPO3(2), respectively. An appreciable augmented regularity in geometrical features of gene nucleotide sequences was realized for wild-type MYB, NFIB, CRTC1, and MAML2, respectively along with their respective mutated variants i.e., MYB-NFIB and CRTC1-MAML2. The change in FD and LC for WTs of PML, RARA, EWSR1, ATF1, and FLI1 and their mutated oncogenes are displayed in Fig. 2(c). In comparison to PML, a decrement in self-similarity and an increment in heterogeneity was observed for the RARA gene sequence. However, their fusion [PML-RARA (1, 2, 3, and 4)] variants do not significantly disrupt the ordering of the WT gene sequence geometry. Similarly, all three mutated variants of EWSR1-ATF1, and four variants of EWSR1-FLI1, exhibited a homogeneous geometry owing to the fusion of respective comparatively heterogeneous geometry of ATF1 and FLI1 with the regular geometrical gene sequence of EWSR1 (Table S2(a)). In Fig. 2(d), variations for FD and LC are exhibited corresponding to EML4, ALK, BAG4, FGFR1, KIF5B, and MET. A lesser rate of alternating base pairs pattern (high LC) was shown by EML4 while an enhanced statistical self-similarity with lesser heterogeneity in nucleotide sequence patterns was observed for ALK. An ALK-like behavior was also realized for EML4-ALK (1), EML4-ALK (2), and EML4-ALK(3) indicating a decrement in the geometric disordering of EML4 owing to the fusion mutation. A lower variance in alternating base pair patterns (low LC) and order (high FD) was shown for BAG4 while FGFR1 showed comprehensible augmented self-similar characteristics and spatial heterogeneity in the nucleotide sequence. Noticeably, their fusion (BAG4-FGFR1) displayed highly ordered geometry with improved pattern of self-similarity in nucleotide pair distribution although with almost no correlation between the base pairs. Also, in contrast to MET, KIF5B displayed a higher degree of heterogeneity in base pair variations and lower self-similarity and hence, disordered geometry. Nonetheless, their fusion (KIF5B-MET) highly disrupted the order in the geometry of individual WT proto-oncogenes. Also, for MET, point mutation and deletion in CDS (c.3001_3021) do not display any change with respect to the WT. The DNA walks for the fusion cases of BAG4-FGFR1 and KIF5B-MET are displayed in Figs. 4 and 5, respectively.

**Fig. 3.**
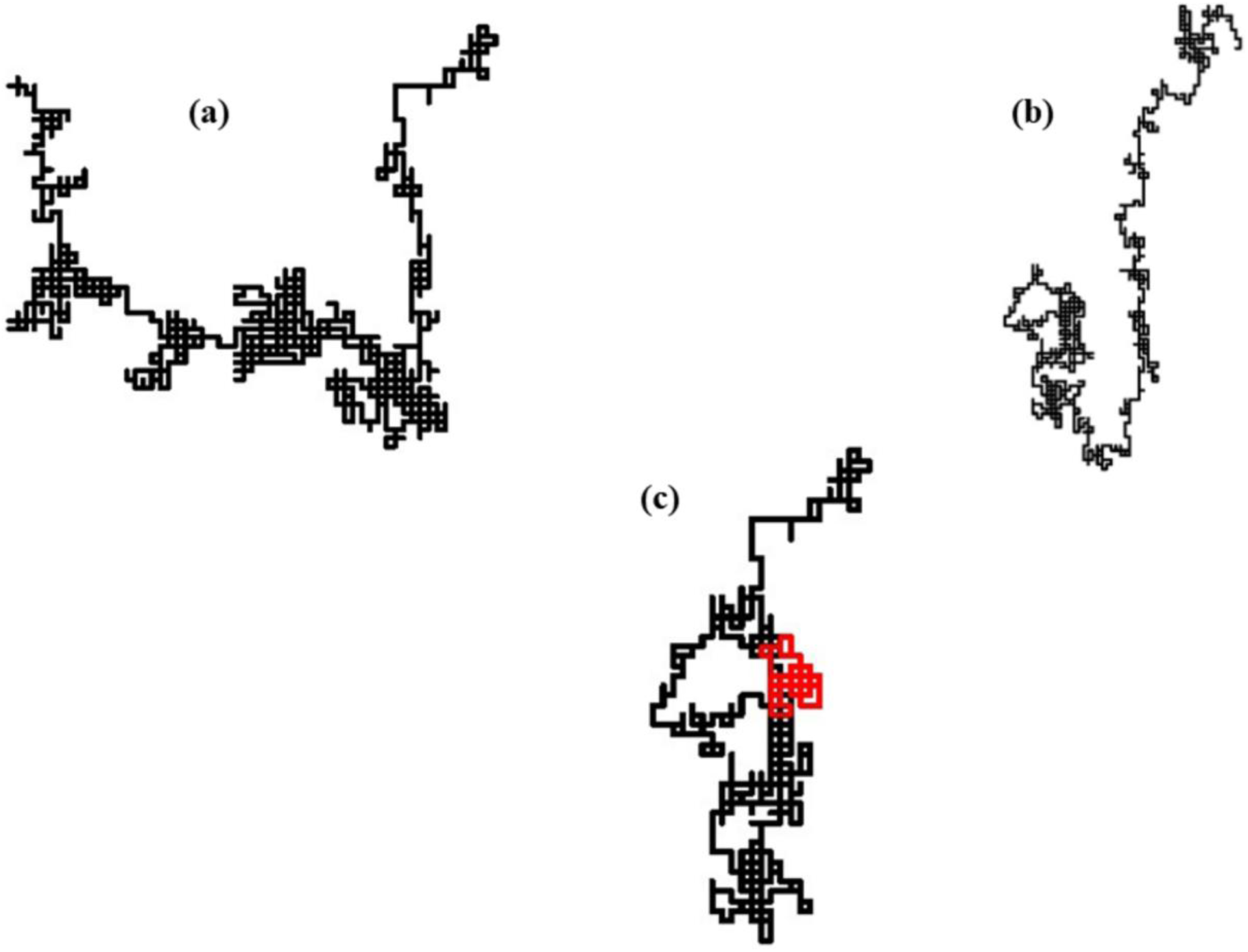
DNA walks corresponding to **(a)** wild-type TMPRSS2, **(b)** wild-type ETV4, and **(c)** fusion variant of TMPRSS2-ETV4.

**Fig. 4.**
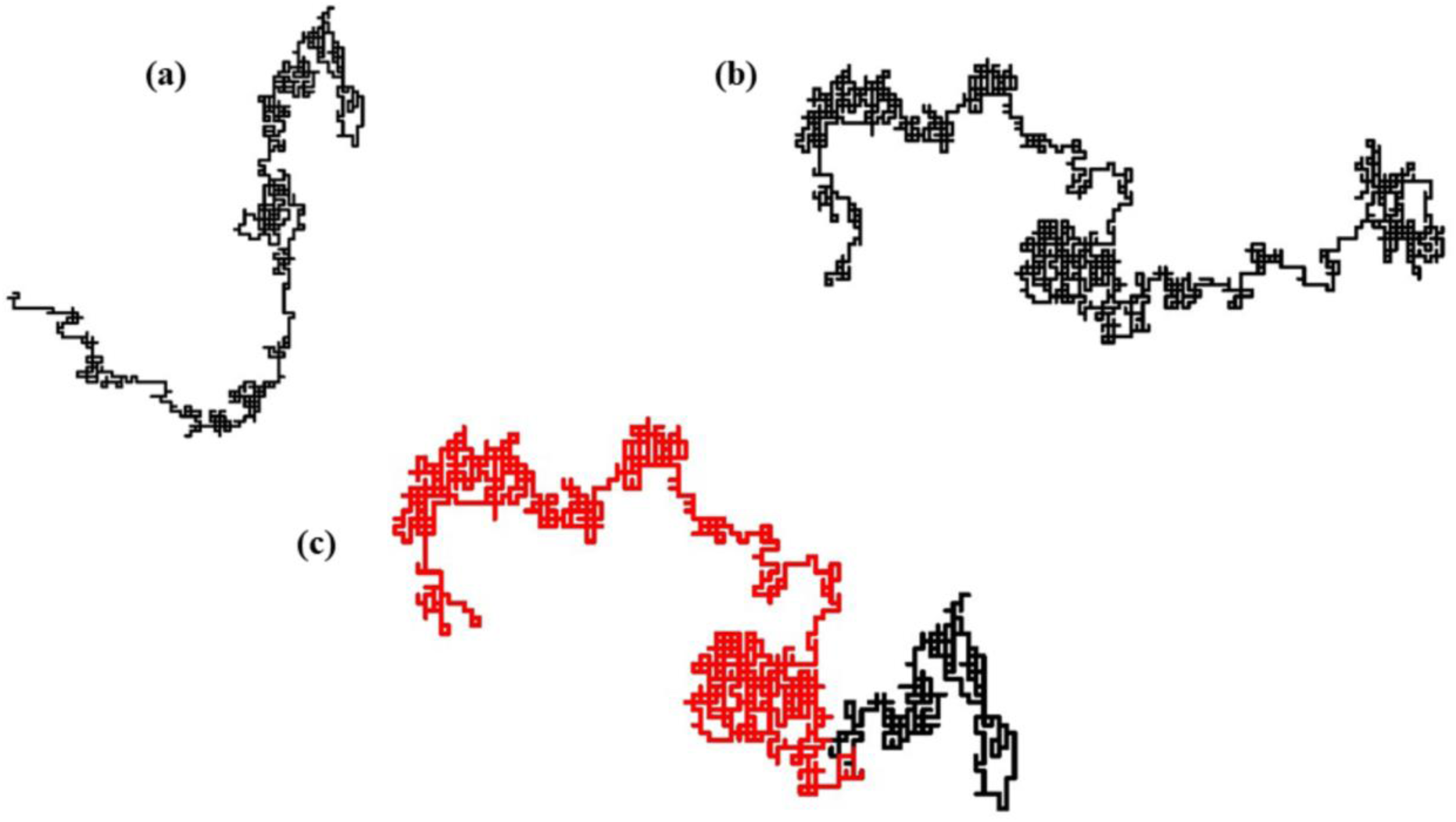
Genetic walks corresponding to **(a)** wild-type BAG4, **(b)** wild-type FGFR1, and **(c)** fusion variant of BAG4-FGFR1.

**Fig. 5.**
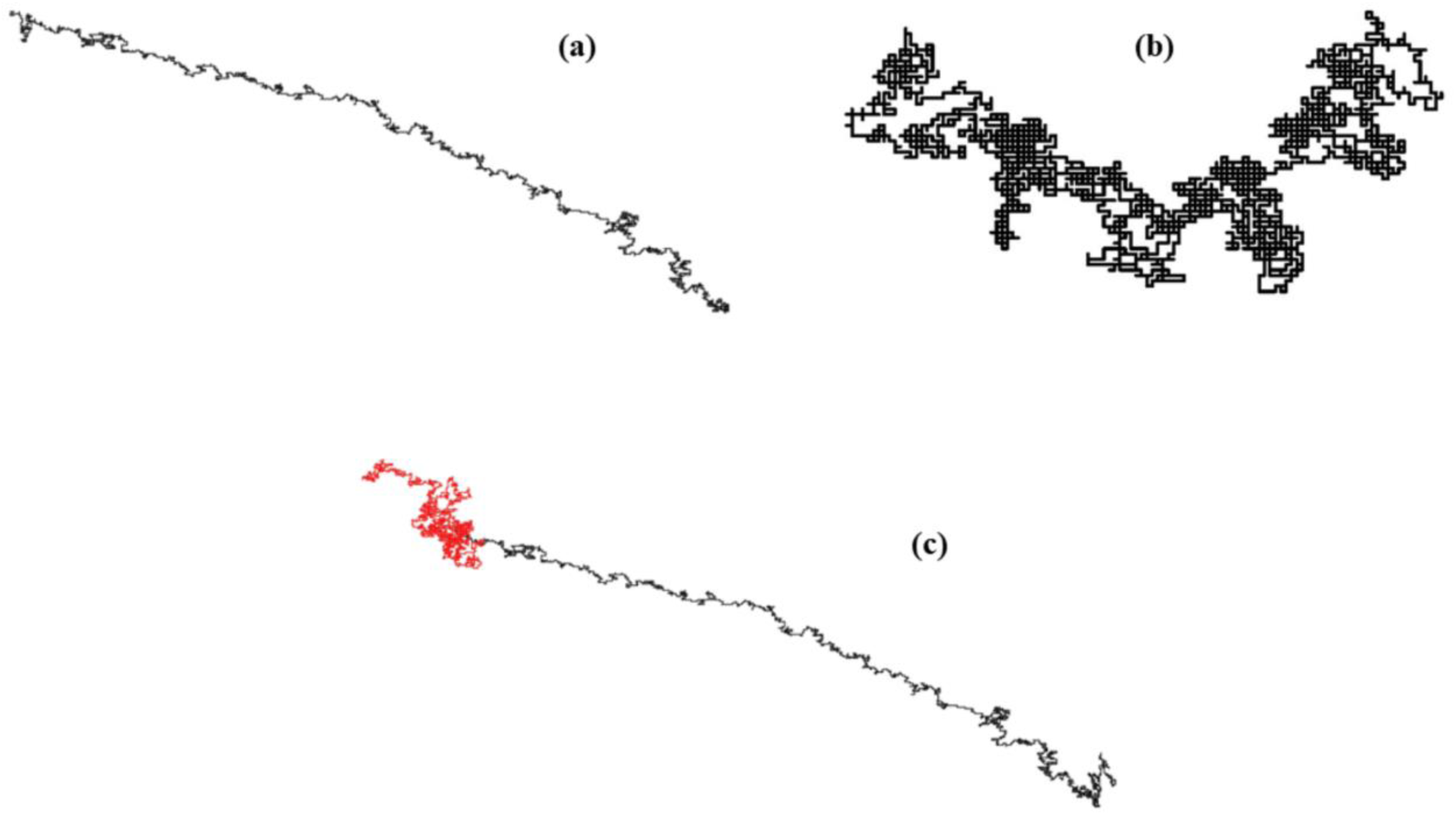
Graphical representation of nucleotide sequence pattern corresponding to **(a)** wild-type KIF5B, **(b)** wild-type MET, and **(c)** KIF5B-MET fusion gene.

The difference in FD and LC measures for WT EGFR, RAD51, fusion gene, point mutations, DEL, and INS corresponding to EGFR are shown in Fig. 6(a). Here, EGFR displayed a notable self-affine characteristic although, with enhancement in heterogeneity subsequently, disordered geometry. On the other hand, a considerable decrement in the rate of alternating base pair patterns, with higher self-affine nucleotide distribution patterns was noted for RAD51. Interestingly, their fusion (EGFR-RAD51) disrupts the geometrical feature in the EGFR with an appreciable increment in the geometric regularity (high FD and low LC). However, almost no change in WT EGFR geometry was realized from the other investigated point, insertion, and deletion mutations. A similar case was also observed for ESR1 where the studied point mutations did not bring any noteworthy change in the WT genes which exhibited a high (low) self-similarity (heterogeneity) in the nucleotide distribution pattern (Fig. 6(b)). The geometrical features were next investigated for KIAA1549 and BRAF WT genes, their fusion gene, along with the point mutated, and DELINS variants of BRAF. The change in FD and LC corresponding to them is shown in Fig. 6(c). A comparatively lower degree of statistical self-similarity and higher spatial heterogeneity in the variation of base pairs was realized in KIAA1549 compared to BRAF. Nonetheless, geometrical disruption was observed in BRAF geometry in the case of fusion (KIAA1549-BRAF) displaying lower regularity and higher heterogeneity in its geometry with long-range correlation between nucleotides. The corresponding DNA walks are shown in Fig. 7. Nonetheless, no significant variations in geometric characteristics were observed for the point mutated, and DELINS variants of BRAF. Finally, we studied the CSD sequence of WT tumour suppressor TP53 gene and its polymorphic forms with point mutations, respectively and scatter plotted the variation in FD and LC as shown in Fig. 6(d). The WT displayed a moderate degree of self-affinity and heterogeneity in the variation of nucleotide patterns. A similar trend in gene nucleotide sequence spatial geometry was realized for all its mutated versions.

**Fig. 6.**
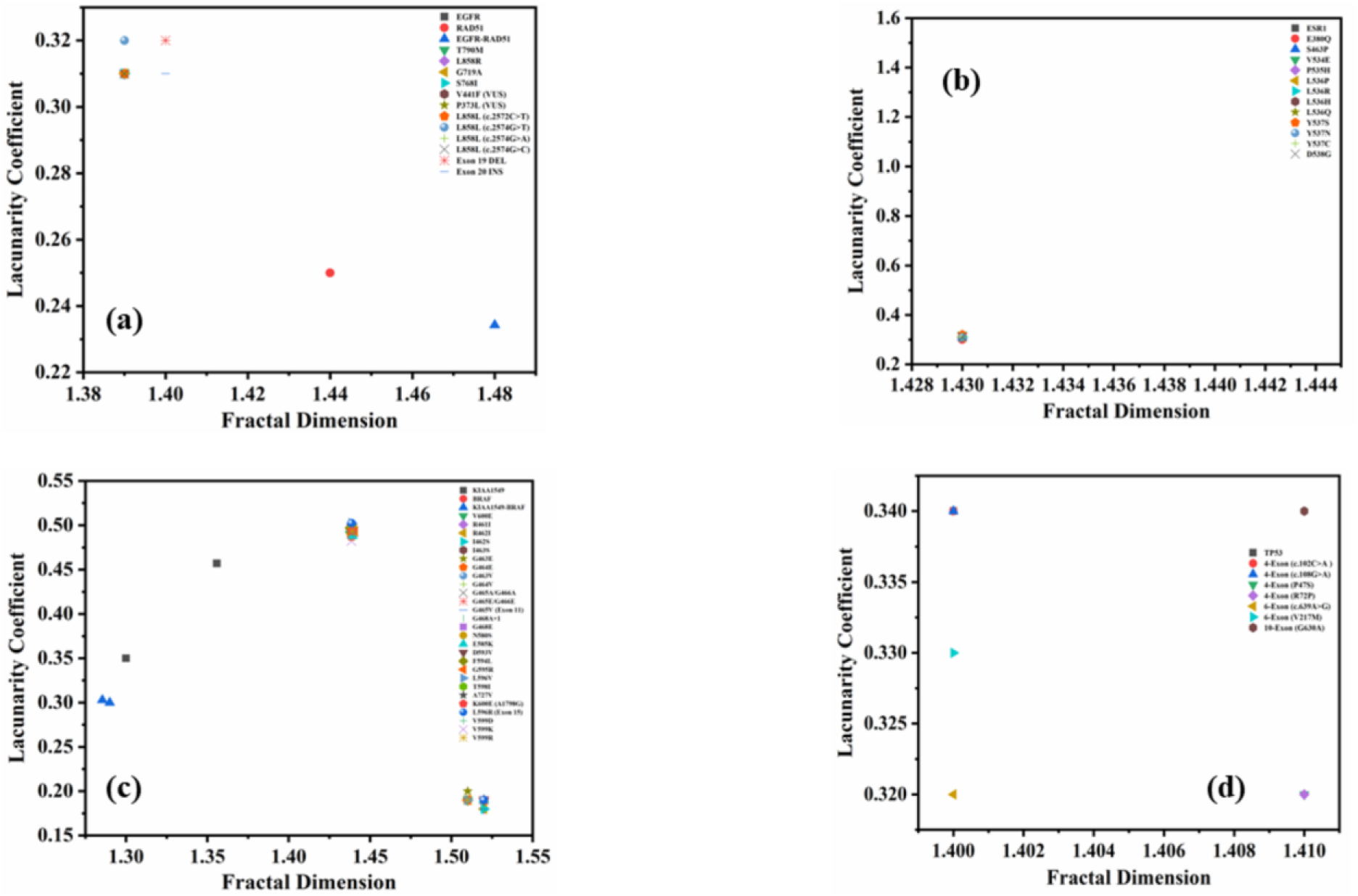
Variation in fractal dimension and lacunarity coefficient in **(a)** wild-type EGFR and RAD51, their fusion form and analyzed mutations in EGFR, **(b)** wild-type ESR1 along with its point mutations, **(c)** wild-type KIAA1549 and BRAF, their fused form, and point mutations in BRAF, and **(d)** wild-type TP53 along with its investigated point mutations.

**Fig. 7.**
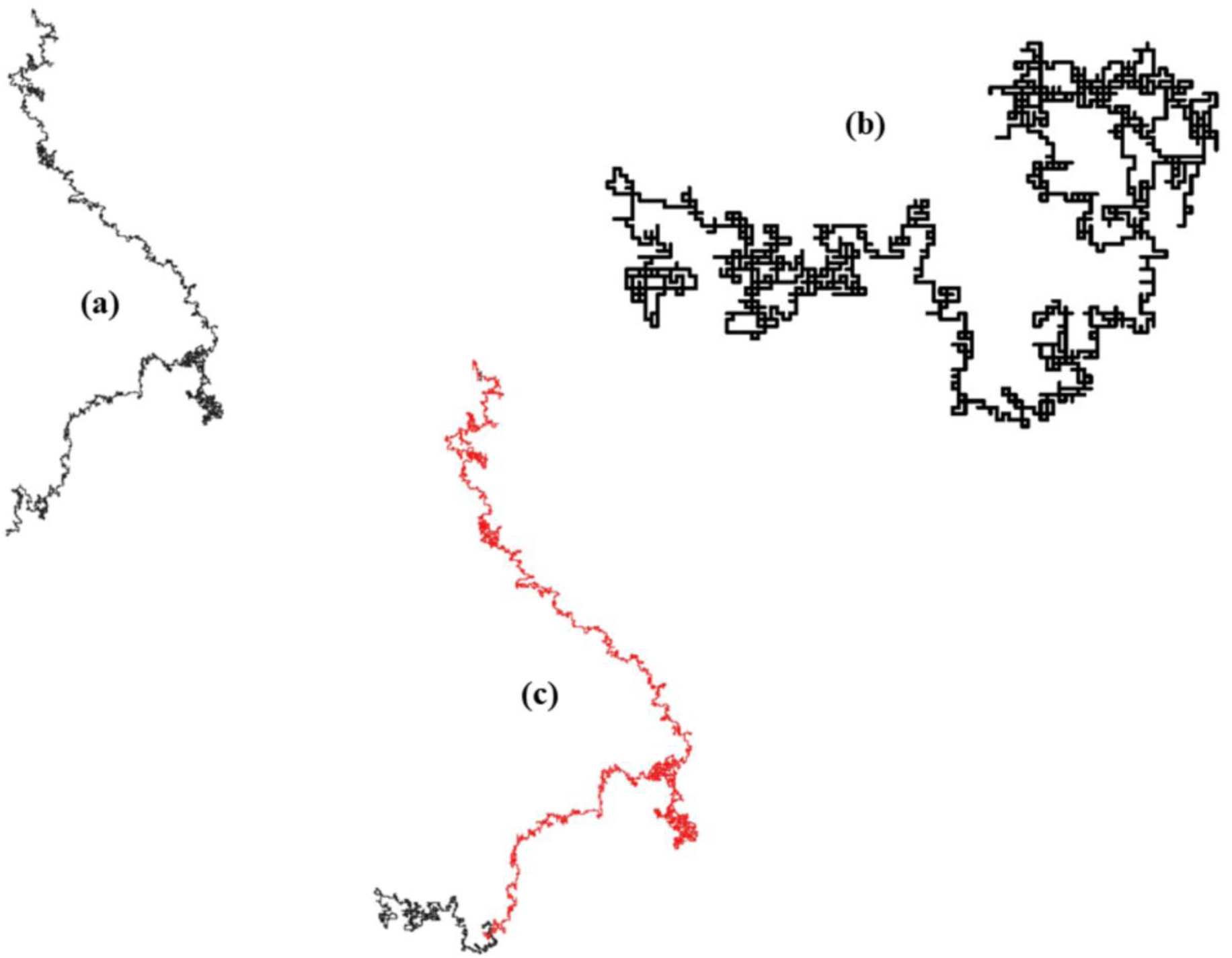
DNA walks for **(a)** wild-type KIAA1549, **(b)** wild-type BRAF, and **(c)** KIAA1549-BRAF fused genes, respectively.

### 3.2 Multi-fractal analysis

#### 3.2.1 DNA walks of wild-type genes, mutated oncogenes, and tumour-suppressor gene

In the present study, we have considered the nucleotide sequences in WT genes and their mutated oncogenic variants as complex systems. With regard to the relation between Hurst exponent and FD, and considering gene sequences as a time series or geometric structure, the mono-fractal parameters (H or FD) signify the scaling of correlation and complexity/self-similarity in them. However, previously, it has been reported in the literature related to complex systems (28–30) that only a single scaling exponent may not be able to comprehensively characterize the present complexity in spatial and temporal patterns. Consequently, a range of scaling exponents (multi-fractality) is required for a comprehensive description of the complexity of the system. Also, multi-fractality has been reported to be absent, vaguely present, and clearly present in random, small-world, and scale-free patterns and networks, respectively (31). The corollary follows that multi-fractal signatures should not be present in case of a random behavior of gene sequences. Therefore, we performed the 2D multifractal analysis of the nucleotide sequences from the DNA walk representation using image analysis and analyzed their multi-fractal behavior in regard to their geometry. The mathematical details regarding the computation of parameters have been described previously (25).

The basic parameters of generalized FD or Renyi dimensions *D_q_* for *q* = 0, 1, and 2 are the capacity, information, and correlation dimensions, respectively. In the case of a multi-fractal system, *D_q_* decreases progressively with an increase in *q* (moment) following the trend *D_0_ > D_1_ > D_2_* with a sigmoidal behavior around *D_0_*. The computed values and trend of the mentioned dimensions were realized to signify the presence of multi-fractal behavior in the investigated DNA gene sequences.

An additional parameter in multi-fractal analysis is the singularity spectrum τ(q) where singularity means irregular or heterogeneous regions in a signal or pattern. In scientific literature, it is repeatedly dictated as the mass exponent owing to its relationship with the Hölder exponent (α) and moment (q). In multi-fractal analysis, a non-linear decrease in mass exponent with an increase in moment indicates the presence of multifractality (32). The trend of τ(q) with -4 ≤ q ≤ 4, for all the studied gene sequences corresponding to WT sequences and their respective mutated variants exhibited a linear trend (Figs. S1, S2, and S3) thereby indicating the absence of multifractality in the gene sequences. The range for the moment parameter was selected according to the reported literature (30).

The results imply the vague presence of multi-fractal characteristics in gene sequence geometry or spatial pattern. In other words, the geometric self-similarity and/or complexity, correlation, and heterogeneity are well captured by the single scaling exponent or mono-fractal parameters in the investigated walks. These results could aid observations in the previous reports (30,33). Nevertheless, we analyzed the multi-fractal spectrums corresponding to the gene sequences to avoid any possible ambiguity in the interpretation of geometry from mono-fractal analysis.

Quantitative information regarding the multi-fractal nature can also be gained from the multifractal spectrum which shows variation in the singularity spectrum f(α) with the Hölder exponent. It is important to note that the mentioned parameters are all related from the Legendre’s transformation, i.e., f(α) = qα(q) – τ(q). Here, the strength of multifractality is quantified from Δα = α_max_ – α_min_ representing the width of the spectrum. In addition, a homogeneous geometrical pattern is associated with q < 0 while heterogeneous and complex geometry is identified for q > 0 (34).

Although the multi-fractality was vaguely observed, the multi-fractal spectrums displayed the complex nature of DNA gene sequence geometry with values of Hölder exponent corresponding to q > 0. The values of α_max_, α_min_, and Δα for Figs. 8-9 and Fig. 10 are tabulated in Table. S3(a) and S3(b), correspondingly. Fig. 8(a) displays the spectrums for genes mentioned in the figure legend. A small strength in multifractality was observed for CRTC1, MAML2, and their fusion variant from the Δα value. A small increase in multifractality was noted for the G12D and G12V mutant versions in comparison to the wild-type KRAS. The SSX1 and SSX2 WT genes showed high values of Δα while SS18 possessed a comparably small multifractal nature. Interestingly, the fusion variants were noted to possess regularity owing to the presence of SS18. For the PIK3CA gene (Δα = 1.24), no significant variation in multi-fractal nature was noted for any of its point mutations (Fig. 8(b)). Fig. 8(c) shows the spectrum for WT SFPQ, TFE3, TMPRSS2, and ETV4 along with their respective fusion mutation. A greater (small) geometrical complexity was noted for SFPQ (TFE3) while SFPQ-TFE1(1) and (2) exhibited a closer resemblance to TFE3 in terms of the multi-fractality strength. On the other hand, ETV4 exhibited greater geometrical complexity in comparison to TMPRSS2 resulting in an augmentation in irregularities in their fused gene sequence. In the case of PTPRK and RSPO3, they both showed a similar degree of multifractality (Fig. 8(d)) while their mutated versions displayed decrement in pattern heterogeneity. Lastly, smaller values of multi-fractality strength were noted for MYB and NFIB wild-type genes and it pointedly decreases in the case of the fusion gene i.e., MYB-NFIB.

**Fig. 8.**
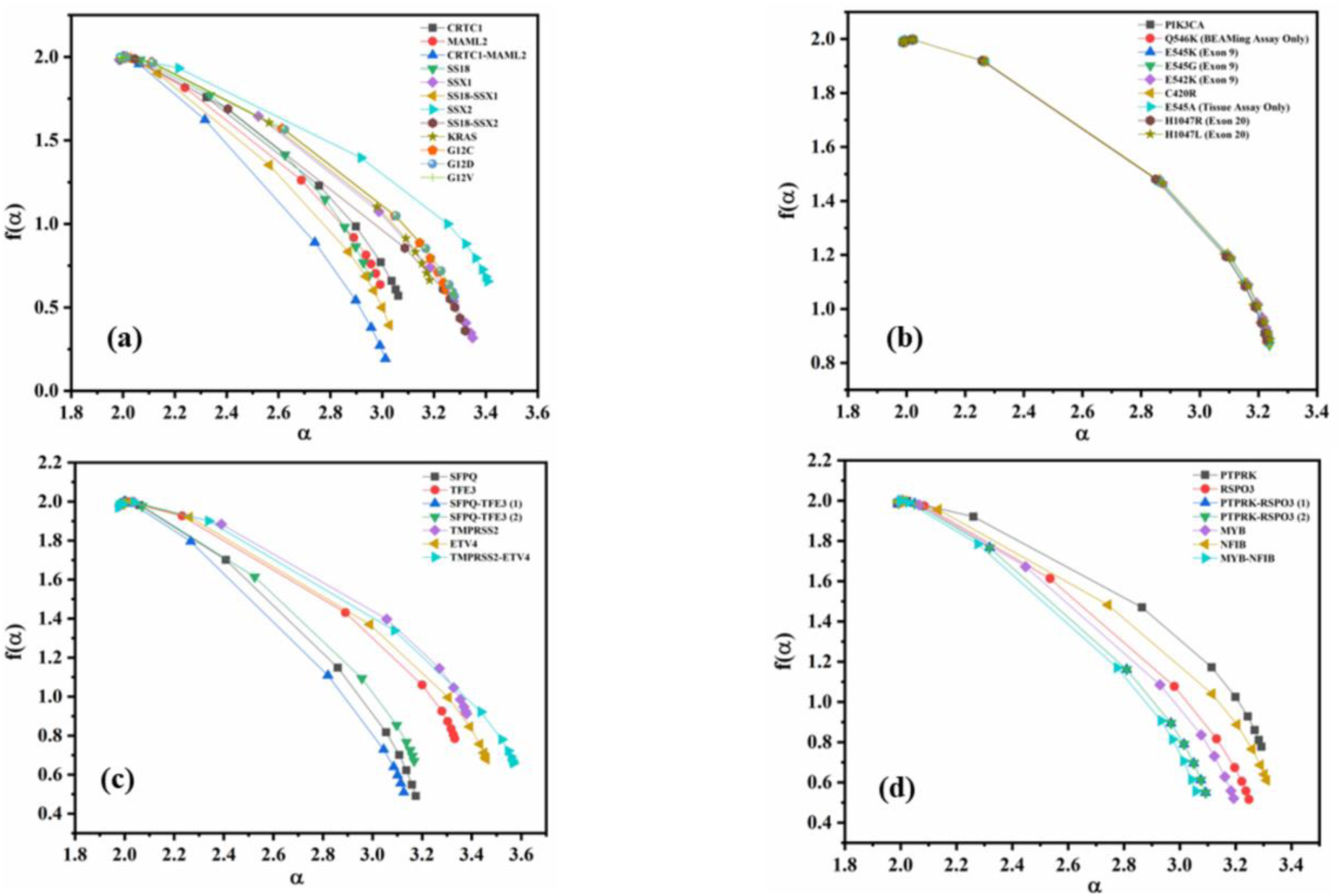
Multi-fractal spectrum between Hölder exponent and singularity spectrum for **(a)** wild-type CRTC1, MAML2, SS18, SSX1, SSX2, and KRAS along with their respective fusion and point mutations, **(b)** wild-type PIK3CA and its point-mutated forms, **(c)** wild-type SFPQ, TFE3, TMPRSS2, and ETV4 with their respective fusion variant, and **(d)** wild-type PTPRK, RSPO3, MYB, and NFIB with their particular fused forms.

**Fig. 9.**
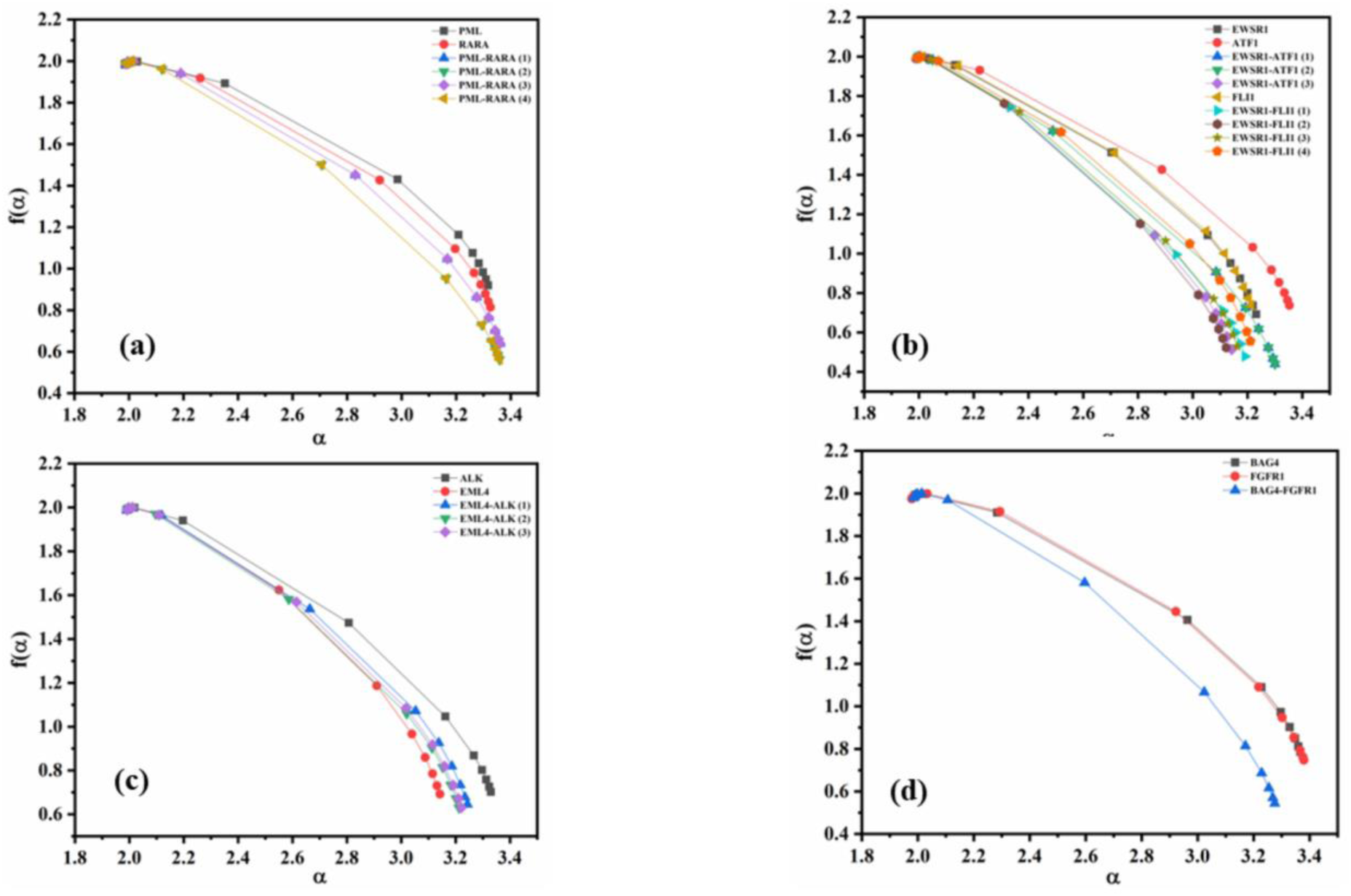
Multi-fractal spectrum for **(a)** wild-type PML and RARA along with their respective fusion form, **(b)** wild-type EWSR1, ATF1, and FLI1 along with individual fusion mutation, **(c)** wild-type ALK and EML4 along with their respective fusion form, and **(d)** wild-type BAG4 and FGFR1 and their fusion form.

**Fig. 10.**
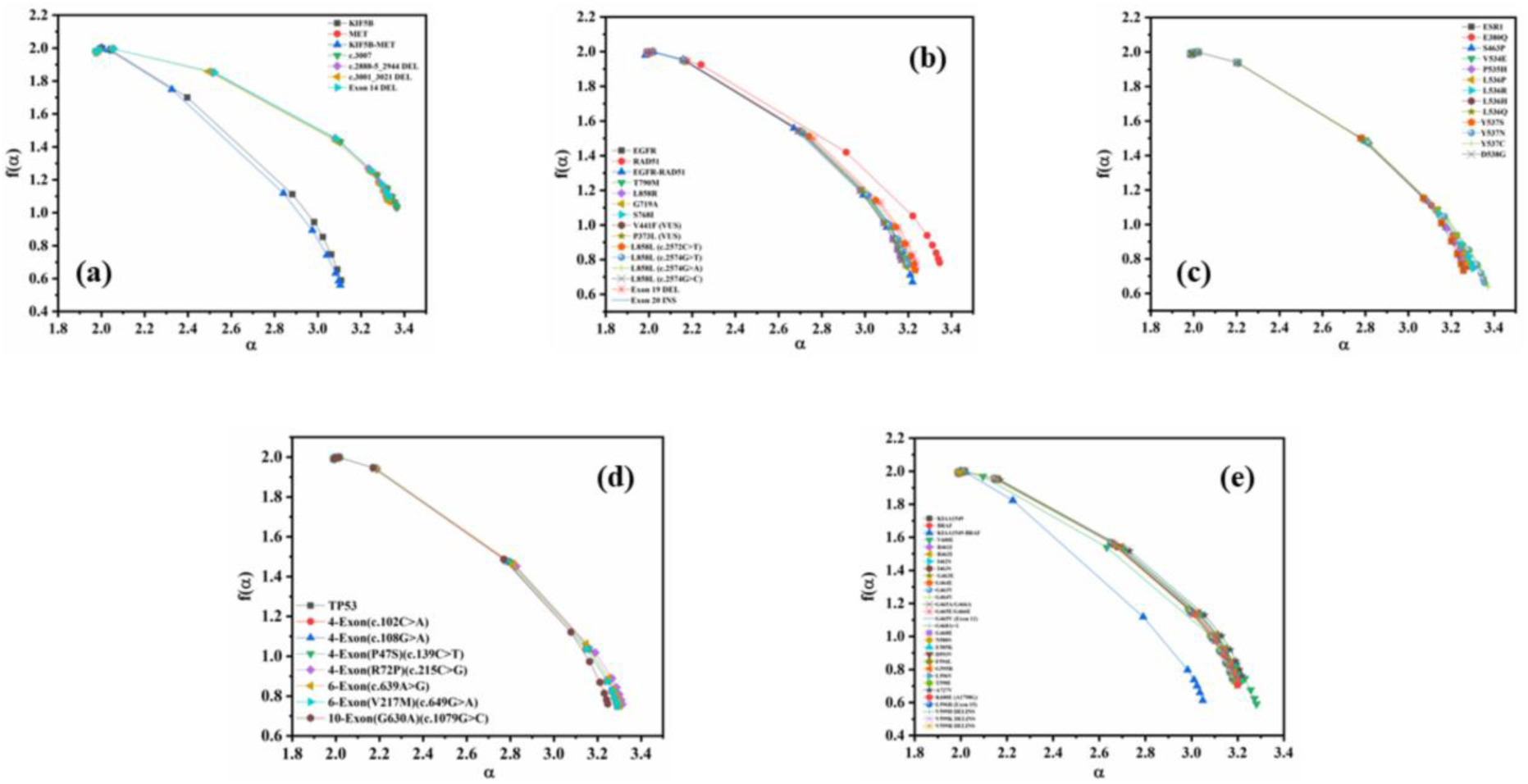
Multi-fractal spectrum for **(a)** wild-type KIF5B and MET along with their respective fusion, and point/deletion mutation in MET **(b)** wild-type EGFR and RAD51 along with individual fusion mutation and point mutations in EGFR **(c)** wild-type ESR1 along with its point mutations, **(d)** wild-type TP53 and its polymorphic forms, and **(e)** wild-type KIAA1549 and BRAF along with their individual fusion mutation and point/DELINS mutations in BRAF.

Figure 9(a, b, c, and d) exhibits the multi-fractal spectrum corresponding to the wild-type genes of PML and RARA, EWSR1, ATF1, and FLI1, ALK and EML4, and BAG4 and FGFR1, respectively. A similar nature of multifractality was realized for PML and RARA with a negligible increase in the mutated versions (Fig. 9(a)). In Fig. 9(b), greater multifractality was noted for ATF1 while FLI1 and EWSR1 exhibited similar multi-fractal behavior. In addition, the mutated variants of EWSR1 and ATF1 displayed enhancement in geometrical complexity except for the third one with enhanced regularity. Also, the fused forms of EWSR1 and FLI showed enhanced homogeneity in their geometry. Next, in the case of ALK and EML4, the former pointed towards a more irregular geometry while the latter showed enhanced regularity. Meanwhile, the fusion forms exhibited an average geometrical characteristic in comparison to the WT counterparts (Fig. 9(c)). Finally, the multi-fractal spectrum is displayed for BAG4, FGFR1, and their fusion variant in Fig. 9(d). Here, a high and similar degree of geometrical complexity was observed for both the WT sequences, however, an appreciable decrement in pattern complexity was highlighted for BAG4-FGFR1.

Fig. 10(a) displays the multi-fractal spectrum corresponding to WT KIF5B, MET, the fusion mutant, point mutation, and deletion variants of MET. A significant augmented in geometric complexity was noted for MET while enhanced regularity was realized for KIF5B. In addition, a significant reduction in the multifractal signature was highlighted by KIF5B-MET however, the point mutation and deletion forms of MET displayed similar behavior in comparison to the WT counterpart. In Fig. 10(b), a higher degree of irregularity in geometry was noted for RAD51 while the opposite was realized for EGFR. Interestingly, the presence of an average behavior in geometry was comprehended from the spectrum width for EGFR-RAD51. Nonetheless, no significant difference in multi-fractal characteristics, in comparison to WT EGFR was observed for the point, deletion, and insertion mutations, respectively. Furthermore, no substantial variation was realized in gene sequence multifractality of wild-type ESR1 and TP53 and their point mutated variants as shown in Fig. 10(c) and 10(d), respectively. Lastly, a decrement in geometric complexity was noted for KIAA1549-BRAF in comparison to the respective WT forms along with point mutated and DELINS forms of BRAF oncogenes in Fig. 10(e).

The results above largely complement the observations from mono-fractal analysis. However, they do not contribute any significant additional information regarding the geometry of investigated gene sequences, thus, strongly highlighting the presence of mono-fractal scaling in the studied sequences. In addition, although we only utilized 2D multi-fractal analysis without any regard to possible trends in gene sequences, the result should hold utilizing techniques like 2D multi-fractal detrended fluctuation analysis (MFDFA) owing to the linear behavior of the mass exponent as reported in earlier work (30).

## 4. Discussion

The concept of symmetry-breaking regarding the combinatorial, geometric, and functional features in cellular components has been argued to describe the notion of complexity and heterogeneity in a system, e.g., cancer (1). Nevertheless, in consideration of the limitation of in investigation of symmetry and symmetry-breaking of combinatorial, geometric, and functional features in complex biological systems, in this study, we specifically focussed on the geometrical features (self-similarity and heterogeneity) highlighting the presence or absence of orderliness in spatial geometry of DNA nucleotide sequences in WT and mutated oncogenes and tumour-suppressor genes relevant to diverse types of cancer. This approach can highlight the qualitative presence or absence of symmetry in the geometrical features in systems of interest.

Here, in regards to DNA gene nucleotide sequences geometry, the presence of order and disorder geometric states facilitated by different types of mutations, i.e., fusion, point mutation, insertion, deletion, and deletion-insertion, was emphasized in the framework of fractal geometry via quantification of mono-fractal (FD and LC) and multi-fractal measures (generalized dimensions, mass exponent, and multi-fractal spectrum), respectively from image analysis of their walks in a 2D space. The approach is also favourable for identifying the type of spatial correlations in the nucleotide base pairs sequencing pattern, if any exist

Due to the large size of the investigated dataset, the main theme of the present work, and the obtained results, the discussion will only be focussed on sequences exhibiting distinct order and disorder geometric states with regard to the mono-fractal measures.

Prostate cancer is generally held to be a localized cancer and exhibits aggressiveness only in a small population of patients. However, it has been reported as the second largest cause of death in males due to its high prevalence in elderly populations (35). Additionally, in over half of prostate cancer patients, the tumour exhibits a recurring translocation or fluctuation, which relocates a gene from the ETS family (such as ERG, ETV1, or ETV4) downstream of a prostate-active gene promoter, leading to the abnormal overexpression of the respective ETS gene (36,37). In particular, the proliferation of prostate cancer cells and their progression through the cell cycle have been reported to be accelerated by the ETV4 gene (38). Also, TMPRSS2 has been characterized as over expressive and a responder to androgen in the prostate. Tomlins *et al*., (39,40) identified and validated the unique repetitions in gene sequence arrangements in a majority of prostate cancer cases via fusion of TMPRSS2 and ETS family members. They hypothesized that dysregulation in gene sequences of the ETS family (e.g. ETV4) via fusion with TMPRSS2 may act as an initiator in the development of prostate cancer (40). In this study, the fusion of TMPRSS2 with ETV4 resulted in an augmentation in self-similar characteristics and a decrement in spatial heterogeneity subsequently enhancing the geometric ordering of the resulting nucleotide sequence in comparison to the WT sequence as quantified by the FD and LC measurements, respectively. This observation can be considered as a geometric analogy to the observed repetitions in gene sequences in the case of TMPRSS2-ETV4 fusion via possible repetition of motifs or sub-sequences and low variance in nucleotide distribution in the fused oncogene.

In many cancers, a significant role is played by the tyrosine kinase receptors known as Fibroblast growth factor receptors (FGFR). They have been reported to dimerize and stimulate intracellular signalling trails accountable for the proliferation and survival of cells following the binding of growth factors of the FGF family (41). In addition, they are involved in the regulation of developmental processes like the initiation of organo- and morpho-genesis, and homeostatic procedures in mature tissues like repair and remodelling (42). Furthermore, the development of a diverse range of novel agents along with improvement in diagnostic tests have been greatly supported by the growing therapeutic relevance of fluctuations in FGFR including fusion (43). In the context of aberrations, mutations are less encountered as compared to amplifications accounting for nearly 26% of tumours altered by FGFR. However, fusions of FGFR have been reported in several types of tumours although, with low incidence, and are classified into two categories, i.e., Type-I and Type-II. In Type-I (FGFR is the 3’ fusion partner), the extracellular and transmembrane domains are excluded from the resulting fusion protein including only the FGFR kinase domain linked to the 5’ partner (44). On the other hand, in Type-II (FGFR is the 5’ fusion partner) the extracellular, transmembrane, and kinase domains remain intact with breakpoint usually occurring at exons 17, 18, or 19 (45). In regards to mutations, this study investigated the presence of geometric order or disorder for 5’-3’ FGFR1 fusion with BAG4, reported in NSCLC (46). It was observed that the self-similar characteristic of the BAG4 nucleotide sequence was highly disrupted in the fusion with FGFR1 along with the display of augmented geometrical order in gene sequence patterns in the fused oncogene. Also, the fused gene displayed comparable self-similar characteristics in nucleotide pattern to FGFR1, as quantified by the FD measure, and this observation could qualitatively aid in explaining the limitation in the detection of rearrangement of FGFR1 gene sequence in the case of BAG4-FGFR1 fusion as reported in another literature (47).

Gene fusion owing to rearrangements in the structure of chromosomes has been reported to be a major driving force of tumourigenesis (48). In a considerable proportion of patients with non-small cell lung carcinoma, genetic instabilities at multiple levels ranging from simple changes in nucleotides, chromosomal and structural rearrangements, gene amplifications, and gain/loss of entire chromosomes are present within the tumours (8). In recent times, the fusion of Kinesin family member 5B (KIF5B) with MET has been discovered in patients with lung adenocarcinoma (49) containing a chimeric fusion of 1-24 and 14-21 or 15-21 exons of KIF5B and MET, respectively (50,51). The investigated mutation in this work was from a fusion of KIF5B and 14-21 exons of MET. It has been observed that oncogenic mutations in the 14-exon splice site of MET can lead to skipping of it consequently, leading to impaired Casitas B-cell lymphoma family E3 ubiquitin ligase binding and decrement in degradation of MET (52). Noticeably, the rearrangement of genes in KIF5B-MET (exon 24-exon 14) results in strong anti-tumour activity of crizotinib in patients with NSCLC (53). We have studied the geometric feature of nucleotide patterns in the KIF5B-MET fusion gene via the DNA walk representation. The results displayed the augmentation in spatial heterogeneity or rate of base pair alterations along with long-range correlation in the resultant nucleotide sequence in contrast to the wild-type MET. In addition, the self-similarity in the base pairs pattern of MET was realized to get highly disrupted owing to the fusion with a comparatively more heterogeneous nucleotide sequence pattern of KIF5B. Furthermore, the investigated c.2888-5_2944 and Exon 14 MET split site mutations displayed comparably high self-similarity and complexity from the wild-type MET. These results could aid in the possible prediction of unidentified somatic alterations indicating the skipping of exon 14 in MET genes as discussed in the reported literature (10).

Brain tumours, especially ones originating from glia of white and grey matter, can harbour gene alterations in v-Raf murine sarcoma viral oncogene homolog B1 (BRAF) (54). Also, its initiation in neural stem and progenitor cells not only endorses the growth of tumours but, subsequently causes oncogene-induced senescence in some low-grade brain tumours (55). In addition, alterations in the BRAF gene have been found in diffusively budding tumours which is connected to poor prognosis (54). In regards to fusion and point mutation-associated complications in brain tumours (pilocytic astrocytomas), the most common is with KIAA1549 and BRAF V600E (56), respectively. Interestingly, the presence of V600E mutation has been reported to be mutually exclusive to that of KIAA1549-BRAF in the case of pilocytic astrocytomas in literature (54). In our study, it was observed that the geometry of the fusion nucleotide sequence presented a decrement in regularity via self-similarity and long-range correlation with high heterogeneity as compared to the wild-type BRAF though, the same geometric features as observed for the WT (ordered state) was also realized for BRAF V600E point mutation (Table S2(b)). The observed distinction possibly highlights different mechanistic variations associated with these mutations and could be considered to investigate the reported difference in prognostic impact from these mutations in several brain tumours (54).

In general, the results from this study support the presence of long-range correlation, mono-fractal scaling, and non-randomness in its gene nucleotide sequences, as reported in previous literature (10,30,57). However, short-range correlation in nucleotide base pair patterns was also realized for some wild-type genes. In the context of geometric order and disorder states, fusion mutations were observed to be of significant interest only since specific fused oncogenes revealed distinct self-similarity and heterogeneity in nucleotide sequence patterns in comparison to their WTs. However, the method (DNA walk) itself might not be able to capture geometrical features due to point mutations. The reason for this is that point mutations involve changes at single nucleotide positions within the DNA gene sequence. These mutations can cause subtle but possibly significant alterations in the local nucleotide structural pattern and the global geometry of the DNA. The DNA walk method, which involves mapping the nucleotide sequence to a 2D spatial representation, may not be sensitive enough to detect these minor but critical changes in geometry. Future work could address this limitation by employing fractal analysis of protein structure data, possibly from crystallographic, spectroscopic, or microscopic characterization. This approach would investigate geometrical changes due to protein folding and/or binding sites, which may provide a more subtle understanding of the structural impacts of point mutations. In addition, integrating the concept of geometric order-disorder in gene sequences with ones in protein conformation might aid in identifying aspects of fluctuations (or noise) in biological regulation.

The geometric behaviour observed in WT might not be generic since different levels of isoforms for a particular gene can exist within a single cell, attributed to the inherent stochastic behaviour of cells. In addition, the walk of WTs may not display the same behaviour for every cell, and similar behaviour can also be seen in mutant genes with multiple isoforms due to dynamic alterations in the regulatory biochemical network from post-transcriptional modifications (10).

Nonetheless, this approach can be fruitful in cancer genomics as this may provide valuable information regarding switch fold and inherently disordered proteins. In addition, the approach can aid in identifying coding regions (exon sequences), non-coding regions (intron sequences), and genomic alterations from epigenetic modifications and environmental factors displaying unique geometric order, disorder, and order-disorder states. In addition, analyzing the geometric properties of coding and non-coding parts of gene sequences might provide clues regarding the impact of fluctuations on the biological pathways regulated by those genes, which can ultimately aid in identifying mechanisms responsible for drug resistance as well as response to therapies.

Additionally, this study dealt with only chromosomal DNA. However, extrachromosomal DNA (ecDNA) has been reported to play a significant role in different types of cancers and associated diseases (58). In this regard, fractal analysis of ecDNA walks might shed light on the global behaviour of ecDNA and consequent protein structures. The approach can also be extended to investigate the features of order and disorder in information in the context of Hurst exponent via investigation of gene sequences as stationary or non-stationary time series.

The present study provides a framework for investigation and possible utilization of the geometric features in the investigation of cancer or its component systems. However, for robust results, the length of sequences should be considered in the investigation of patterns in complementary sequences as it could affect the correlation measure. In addition, this kind of study can shed light on the evolution process of DNA and epigenetics. Nonetheless, we believe researchers should perform fractal analysis of functional features (gene-gene and protein-protein interaction networks) in combination with geometric features to gain deeper insights into the complexity of cancer.

Finally, we demonstrate bridging between clinical observations and translational research by an interdisciplinary approach via integrating genetic and geometric analyses. The approach can be employed to understand the complexities of cancer and its outcomes along with possibly contributing to team medicine via designing and conducting innovative research.

## 5. Conclusion

Cancer is a complex adaptive system and therefore, has been viewed for symmetry-breaking where the system’s complexity is embodied by its combinatorial, geometric, and functional features. However, such an approach can have certain limitations in direct interpretations due to a lack of understanding of the physical laws governing symmetry in complex systems. As an alternative, here, we investigated the implication of geometric features (self-similarity and heterogeneity), where their combination highlighted the presence of ordered and disordered geometric states of DNA sequences (WT and their mutated variants) related to various types of cancer in the framework of fractal geometry utilizing the DNA walk representation. FD and LC coefficient quantified the geometrical self-similarity and heterogeneity in the analyzed sequence patterns and identified spatial correlations by the FD measure. Interestingly, distinct fractal geometric features were realized in the case of gene fusions. The 2D multifractal analysis highlighted the presence of a single exponent in the scaling of investigated gene sequence self-similar and heterogeneous behavior. The distinct fractal features were suggested to be qualitatively related, as geometric analogies, to specific observations reported in the literature corresponding to specific cancer types. Finally, the advantages and limitations of the present approach as well as future research directions were highlighted.

## Author Contributions

Conceptualization, R.S. and M.K.J; Methodology, A.D. and M.K.J; Formal analysis, A.D., M.S., A.S., R.G. and M.P; Investigation, A.D., Data curation, M.S., A.S., A.S., R.G. and M.P; Writing-original draft preparation, A.D.; Writing-review and editing, A.D., J.F., I.M., P.K., M.K.J and R.S.

## Data Availability Statement

The data presented in this study are available on request from the corresponding author.

## Conflict of Interest

The authors declare no conflict of interest.

## Acknowledgment

A.D. acknowledges the Department of Biotechnology (DBT)-India for the Research Associate fellowship vide Award Letter No. DBT-RA/2023/January/NE/3594.

## Notes

### Competing Interest Statement

The authors have declared no competing interest.

## References

1. Frost JJ, Pienta KJ, Coffey DS. Symmetry and symmetry breaking in cancer: a foundational approach to the cancer problem. Oncotarget. 2018;9(14):11429–40.

2. Lucia U, Grisolia G, Deisboeck TS. A non-equilibrium thermodynamic approach to symmetry breaking in cancer. AAPP Atti della Accad Peloritana dei Pericolanti, Cl di Sci Fis Mat e Nat. 2021;99(1):1–12.

3. Weinberg RA. Coming full circle - From endless complexity to simplicity and back again. Cell. 2014;157(1):267–71.

4. Longo G, Montévil M. From physics to biology by extending criticality and symmetry breakings. Prog Biophys Mol Biol. 2011;106(2):340–7.

5. Longo G, Montévil M. Perspectives on Organisms: Biological Time, Symmetries and Singularities. Springer-Verlag Berlin Heidelberg. 2013.

6. Coffey DS. Self-organization, complexity and chaos: The new biology for medicine. Nat Med. 1998;4(8):882–85.

7. Vogelstein B, Kinzler KW. Cancer genes and the pathways they control. Nat Med. 2004;10(8):789–99.

8. Shen Z. Genomic instability and cancer: An introduction. J. Mol. Cell Biol. 2011;3(1):1– 3.

9. Hanahan D. Hallmarks of Cancer: New Dimensions. Cancer Discov. 2022;12(1):31–46.

10. Hewelt B, Li H, Jolly MK, Kulkarni P, Mambetsariev I, Salgia R. The DNA walk and its demonstration of deterministic chaos - Relevance to genomic alterations in lung cancer. Bioinformatics. 2019;35(16):2738–48

11. Herbst RS, Morgensztern D, Boshoff C. The biology and management of non-small cell lung cancer. Nature. 2018;553:446–54.

12. Faguet GB. A brief history of cancer: Age-old milestones underlying our current knowledge database. Int. J. Cancer 136(2014);2022–36.

13. Lyons SM, Alizadeh E, Mannheimer J, Schuamberg K, Castle J, Schroder B, et al. Changes in cell shape are correlated with metastatic potential in murine and human osteosarcomas. Biology Open. 2016;5:289–99.

14. Lennon FE, Cianci GC, Cipriani NA, Hensing TA, Zhang HJ, Chen CT, et al. Lung cancer-a fractal viewpoint. Nat. Rev. Clin. Oncol. 2015;12:664–75.

15. Metze K. Fractal dimension of chromatin: Potential molecular diagnostic applications for cancer prognosis. Expert Rev. Mol. Diagn. 2013;13(7):719–35.

16. Sokolov I. Fractals: a possible new path to diagnose and cure cancer ? Future Oncol. 2015;11:3049–51.

17. d’Onofrio A. Fractal growth of tumors and other cellular populations: Linking the mechanistic to the phenomenological modeling and vice versa. Chaos, Solit. Fractals. 2009;41(2):875–80.

18. Verma G, Luciani ML, Palombo A, Metaxa L, Panzironi G, Pediconi F, et al. Microcalcification morphological descriptors and parenchyma fractal dimension hierarchically interact in breast cancer: A diagnostic perspective. Comput. Biol. Med. 2018;93:1–6.

19. Karlin S, Brendel V. Patchiness and correlations in DNA sequences. Science. 1993;259:677–80.

20. Namazi H, Kulish VV., Delaviz F, Delaviz A. Diagnosis of skin cancer by correlation and complexity analyses of damaged DNA. Oncotarget. 2015;6(40):42623–31.

21. Namazi H, Kiminezhadmalaie M. Diagnosis of Lung Cancer by Fractal Analysis of Damaged DNA. Comput. Math. Methods Med. 2015;1–11.

22. Shahriyari L, Komarova NL. Symmetric vs. asymmetric stem cell divisions: An Adaptation against cancer? PLoS One. 2013;8(10):e76195.

23. Buldyrev SV. Power Law Correlations in DNA. Madame Curie Bioscience Database. 2023;1–37. https://www.ncbi.nlm.nih.gov/books/NBK6262/

24. Das A, Jaiswal J, Yadav RP, Mittal AK, Ţălu Ş, Kumar S. Complex roughening dynamics and wettability mechanism in MoS_2_ thin films — A system theoretic approach. Phys. A. 2023;624:128989.

25. Torre IG, Heck RJ, Tarquis AM. MULTIFRAC: An ImageJ plugin for multiscale characterization of 2D and 3D stack images. SoftwareX. 2020;12:100574.

26. Mandelbrot BB. The Fractal Geometry of Nature. W.H.Freeman and Co. 1982.

27. Abramson G, Cerdeira HA, Bruschi C. Fractal properties of DNA walks. BioSystems. 1999;49:63–70.

28. Das A, Jaiswal J, Borah CK, Ruti I, Matos RS, Pinto EP, et al. Correlating the Nonlinear Roughening and Optical Properties of Anatase Thin Films — A Fractal Geometric Approach. Adv. Thoery Simul. 2023;6(9):2300238.

29. Torre IG, Losada JC, Tarquis AM. Multiscaling properties of soil images. Biosyst. Eng. 2018;168:133–41.

30. Rosas A, Nogueira E, Fontanari JF. Multifractal analysis of DNA walks and trails. Phys.Rev. E 2002;66:061906.

31. Liu JL, Yu ZG, Anh V. Determination of multifractal dimensions of complex networks by means of the sandbox algorithm. Chaos. 2015;25:023103.

32. Das A, Yadav RP, Chawla V, Kumar S, Ţălu Ş, Pinto EP, et al. Analyzing the surface dynamics of titanium thin films using fractal and multifractal geometry. Mater. Today Commun. 2021;27:102385.

33. Arneodo A, Bacry E, Graves PV., Muzy JF. Characterizing long-range correlations in DNA sequences from wavelet analysis. Phys. Rev. Lett. 1995;74(16):3293–96.

34. Ţălu Ş, Morozov IA, Yadav RP. Multifractal analysis of sputtered indium tin oxide thin film surfaces. Appl. Surf. Sci. 2019;484:892–8.

35. Siegel RL, Miller KD, Jemal A. Cancer statistics, 2019. CA Cancer J. Clin. 2019:69:7– 34.

36. Arora K, Barbieri CE. Molecular Subtypes of Prostate Cancer. Curr. Oncol. Rep. 2018;20:58.

37. Lin C, Salzillo TC, Bader DA, Wilkenfeld SR, Awad D, Pulliam TL, et al. Prostate Cancer Energetics and Biosynthesis. Advances in Experimental Medicine and Biology. 2019;185–237.

38. Cosi I, Pellecchia A, De Lorenzo E, Torre E, Sica M, Nesi G, et al. ETV4 promotes late development of prostatic intraepithelial neoplasia and cell proliferation through direct and p53-mediated downregulation of p21. J. Hematol. Oncol. 2020;13:112.

39. Tomlins S a, Rhodes DR, Perner S, Dhanasekaran SM, Mehra R, Sun X-W, et al. Recurrent Fusion of TMPRSS2 and ETS Transcription Factor Genes in Prostate Cancer. Science. 2005;310:644–8.

40. Tomlins SA, Mehra R, Rhodes DR, Smith LR, Roulston D, Helgeson BE, et al. TMPRSS2:ETV4 gene fusions define a third molecular subtype of prostate cancer. Cancer Res. 2006;66(7):3396–400.

41. Babina IS, Turner NC. Advances and challenges in targeting FGFR signalling in cancer. Nat. Rev. Cancer. 2017;17(5):318–32.

42. Goetz R, Mohammadi M. Exploring mechanisms of FGF signalling through the lens of structural biology. Nat. Rev. Mol. Cell Biol. 2013;14(3):166–80.

43. De Luca A, Abate RE, Rachiglio AM, Maiello MR, Esposito C, Schettino C, et al. FGFR fusions in cancer: From diagnostic approaches to therapeutic intervention. Int. J. Mol. Sci. 2020;21(18):1–18.

44. Helsten T, Elkin S, Arthur E, Tomson BN, Carter J, Kurzrock R. The FGFR landscape in cancer: Analysis of 4,853 tumors by next-generation sequencing. Clin. Cancer Res. 2016;22(1):259–67.

45. Gallo LH, Nelson KN, Meyer AN, Donoghue DJ. Functions of Fibroblast Growth Factor Receptors in cancer defined by novel translocations and mutations. Cytokine Growth Factor Rev. 2015;26:425–49.

46. Qin A, Johnson A, Ross JS, Miller VA, Ali SM, Schrock AB, et al. Detection of Known and Novel FGFR Fusions in Non–Small Cell Lung Cancer by Comprehensive Genomic Profiling. J. Thorac. Oncol. 2018;14(1):54–62.

47. Wang R, Wang L, Li Y, Hu H, Shen L, Shen X, et al. FGFR1/3 tyrosine kinase fusions define a unique molecular subtype of non-small cell lung cancer. Clin. Cancer Res. 2014;20:4107–14.

48. Akbari Moqadam F, Lange-Turenhout EAM, Ariës IM, Pieters R, den Boer ML. MiR-125b, miR-100 and miR-99a co-regulate vincristine resistance in childhood acute lymphoblastic leukemia. Leuk. Res. 2013;37:1315–21.

49. Greulich H. The genomics of lung adenocarcinoma: Opportunities for targeted therapies. Genes & Cancer. 2010;1(12):1200–10.

50. Stransky N, Cerami E, Schalm S, Kim JL, Lengauer C. The landscape of kinase fusions in cancer. Nat. Commun. 2014;5:4846.

51. Plenker D, Bertrand M, de Langen AJ, Riedel R, Lorenz C, Scheel AH, et al. Structural alterations of MET trigger response to MET kinase inhibition in lung adenocarcinoma patients. Clin. Cancer Res. 2018;24:1337–43.

52. Gow CH, Liu YN, Li HY, Hsieh MS, Chang SH, Luo SC, et al. Oncogenic Function of a KIF5B-MET Fusion Variant in Non-Small Cell Lung Cancer. Neoplasia. 2018;20(8):838–47.

53. Cho JH, Ku BM, Sun JM, Lee SH, Ahn JS, Park K, et al. KIF5B-MET Gene Rearrangement with Robust Antitumor Activity in Response to Crizotinib in Lung Adenocarcinoma. J. Thorac. Oncol. 2018;13:e29–31.

54. Behling F, Schittenhelm J. Oncogenic BRAF alterations and their role in brain tumors. Cancers. 2019;11:794.

55. Raabe EH, Lim KS, Kim JM, Meeker A, Mao XG, Nikkhah G, et al. BRAF activation induces transformation and then senescence in human neural stem cells: A pilocytic astrocytoma model. Clin. Cancer Res. 2011;17:3590–9.

56. Yao Z, Torres NM, Tao A, Gao Y, Luo L, Li Q, et al. BRAF Mutants Evade ERKDependent Feedback by Different Mechanisms that Determine Their Sensitivity to Pharmacologic Inhibition. Cancer Cell. 2015;28:370–83.

57. Provata A, Nicolis C, Nicolis G. DNA viewed as an out-of-equilibrium structure. Phys. Rev. E. 2014;89:052105.

58. Zhao Y, Yu L, Zhang S, Su X, Zhou X. Extrachromosomal circular DNA: Current status and future prospects. eLife. 2022;11:e81412.

