## Supplementary Material for "DNA walk of specific fused oncogenes exhibit distinct fractal geometric characteristics in nucleotide patterns"

[†]**Current affiliation**

| Gene | Fractal Dimension | Lacunarity Coefficient |
|---|---|---|
| EML4 | 1.30 | 0.41 |
| EGFR | 1.39 | 0.31 |
| ALK | 1.41 | 0.36 |
| BAG4 | 1.38 | 0.30 |
| FGFR1 | 1.46 | 0.34 |
| RAD51 | 1.44 | 0.25 |
| MET | 1.57 | 0.18 |
| KIF5B | 1.21 | 0.24 |
| PML | 1.40 | 0.18 |
| RARA | 1.32 | 0.25 |
| ATF1 | 1.35 | 0.34 |
| EWSR1 | 1.22 | 0.21 |
| SFPQ | 1.24 | 0.40 |
| TFE3 | 1.35 | 0.25 |
| TMPRSS2 | 1.41 | 0.27 |
| ETV4 | 1.43 | 0.36 |
| PTPRK | 1.41 | 0.34 |
| RSPO3 | 1.29 | 0.57 |
| MYB | 1.27 | 0.28 |
| NFIB | 1.30 | 0.33 |
| BRAF | 1.52 | 0.19 |
| KIAA1549 | 1.30 | 0.35 |
| CRTC1 | 1.22 | 0.22 |
| MAML2 | 1.16 | 0.22 |
| FLI1 | 1.31 | 0.28 |
| SS18 | 1.37 | 0.23 |
| SSX1 | 1.35 | 0.28 |
| SSX2 | 1.34 | 0.27 |
| TP53 | 1.40 | 0.34 |
| ESR1 | 1.43 | 0.31 |
| KRAS | 1.30 | 0.29 |
| PIK3CA | 1.37 | 0.26 |

**Table S1.** Computed fractal dimension and lacunarity coefficient values for the wild-type variant of the studied oncogenes. The values are rounded up to two-significant digits

| Gene | Fractal Dimension | Lacunarity Coefficient |
|---|---|---|
| SS18 | 1.37 | 0.23 |
| SSX1 | 1.35 | 0.28 |
| SS18-SSX1 | 1.36 | 0.27 |
| SSX2 | 1.34 | 0.27 |
| SS18-SSX2 | 1.27 | 0.27 |
| KRAS | 1.30 | 0.29 |
| G12C | 1.30 | 0.29 |
| G12D | 1.30 | 0.29 |
| G12V | 1.30 | 0.29 |
| PIK3CA | 1.37 | 0.26 |
| Q546K (BEAMing Assay Only) | 1.37 | 0.26 |
| E545K (Exon 9) | 1.37 | 0.26 |
| E545G (Exon 9) | 1.37 | 0.26 |
| E542K (Exon 9) | 1.37 | 0.26 |
| C420R | 1.42 | 0.26 |
| E545A (Tissue Assay Only) | 1.42 | 0.26 |
| H1047R (Exon 20) | 1.37 | 0.26 |
| H1047L (Exon 20) | 1.37 | 0.26 |
| SFPQ | 1.24 | 0.40 |
| TFE3 | 1.35 | 0.25 |
| SFPQ-TFE3(1) | 1.27 | 0.42 |
| SFPQ-TFE3(2) | 1.28 | 0.22 |
| TMPRSS2 | 1.41 | 0.27 |
| ETV4 | 1.43 | 0.36 |
| TMPRSS2-ETV4 | 1.48 | 0.22 |
| PTPRK | 1.41 | 0.34 |
| RSPO3 | 1.29 | 0.57 |
| PTPRK-RSPO3(1) | 1.42 | 0.53 |
| PTPRK-RSPO3(2) | 1.42 | 0.53 |
| MYB | 1.27 | 0.28 |
| NFIB | 1.30 | 0.33 |
| MYB-NFIB | 1.22 | 0.25 |
| CRTC1 | 1.22 | 0.22 |
| MAML2 | 1.16 | 0.22 |
| CRTC1-MAML2 | 1.16 | 0.21 |
| PML | 1.40 | 0.18 |
| RARA | 1.32 | 0.25 |
| PML-RARA(1) | 1.38 | 0.24 |
| PML-RARA(2) | 1.37 | 0.23 |
| PML-RARA(3) | 1.38 | 0.24 |
| PML-RARA(4) | 1.37 | 0.23 |
| EWSR1 | 1.22 | 0.21 |
| ATF1 | 1.35 | 0.34 |
| EWSR1-ATF1(1) | 1.22 | 0.22 |
| EWSR1-ATF1(2) | 1.22 | 0.22 |
| EWSR1-ATF1(3) | 1.21 | 0.21 |
| FLI1 | 1.31 | 0.28 |
| EWSR1-FLI1(1) | 1.22 | 0.19 |
| EWSR1-FLI1(2) | 1.23 | 0.19 |
| EWSR1-FLI1(3) | 1.23 | 0.22 |
| EWSR1-FLI1(4) | 1.25 | 0.23 |
| EML4 | 1.34 | 0.44 |
| ALK | 1.40 | 0.33 |
| EML4-ALK(1) | 1.40 | 0.33 |

| | | |
|---|---|---|
| EML4-ALK(2) | 1.37 | 0.35 |
| EML4-ALK(3) | 1.40 | 0.33 |
| BAG4 | 1.38 | 0.30 |
| FGFR1 | 1.46 | 0.34 |
| BAG4-FGFR1 | 1.48 | 0.30 |
| KIF5B | 1.21 | 0.24 |
| MET | 1.57 | 0.18 |
| KIF5B-MET | 1.25 | 0.60 |
| c.3007 | 1.57 | 0.18 |
| c.2888-5_2944 DEL | 1.57 | 0.18 |
| c.3001_3021 DEL | 1.57 | 0.18 |
| Exon 14 DEL | 1.58 | 0.18 |

**Table S2(a).** Computed values of fractal dimension and lacunarity coefficient corresponding to the analyzed DNA walks of respective wild-type and mutated oncogenes in Fig. 2

| Gene | Fractal Dimension | Lacunarity Coefficient |
|---|---|---|
| EGFR | 1.39 | 0.31 |
| RAD51 | 1.44 | 0.25 |
| EGFR-RAD51 | 1.48 | 0.23 |
| T790M | 1.39 | 0.31 |
| L858R | 1.39 | 0.31 |
| G719A | 1.39 | 0.31 |
| S768I | 1.39 | 0.31 |
| V441F (VUS) | 1.39 | 0.31 |
| P373L (VUS) | 1.39 | 0.31 |
| L858L (c.2572C>T) | 1.39 | 0.31 |
| L858L (c.2574G>T) | 1.39 | 0.32 |
| L858L (c.2574G>A) | 1.39 | 0.31 |
| L858L (c.2574G>C) | 1.39 | 0.31 |
| Exon 19 DEL | 1.40 | 0.32 |
| Exon 20 INS | 1.40 | 0.31 |
| ESR1 | 1.43 | 0.31 |
| E380Q | 1.43 | 0.30 |
| S463P | 1.43 | 0.31 |
| V534E | 1.43 | 0.32 |
| P535H | 1.43 | 0.31 |
| L536P | 1.43 | 0.31 |
| L536R | 1.43 | 0.31 |
| L536H | 1.43 | 0.31 |
| L536Q | 1.43 | 0.31 |
| Y537S | 1.43 | 0.32 |
| Y537N | 1.43 | 0.31 |
| Y537C | 1.43 | 0.31 |
| D538G | 1.43 | 0.31 |
| KIAA1549 | 1.30 | 0.35 |
| BRAF | 1.52 | 0.19 |
| KIAA1549-BRAF | 1.29 | 0.30 |
| V600E | 1.51 | 0.19 |
| R461I | 1.52 | 0.18 |
| R462I | 1.52 | 0.18 |
| I462S | 1.52 | 0.18 |
| I463S | 1.51 | 0.19 |
| G463E | 1.51 | 0.20 |
| G464E | 1.52 | 0.19 |
| G463V | 1.52 | 0.19 |
| G464V | 1.52 | 0.18 |
| G465A/G466A | 1.52 | 0.19 |
| G465E/G466E | 1.52 | 0.19 |
| G465V (Exon 11) | 1.52 | 0.18 |
| G468A+1 | 1.52 | 0.19 |
| G468E | 1.52 | 0.19 |
| N580S | 1.52 | 0.19 |
| E585K | 1.52 | 0.19 |
| D593V | 1.52 | 0.19 |
| F594L | 1.51 | 0.19 |
| G595R | 1.51 | 0.19 |
| L596V | 1.51 | 0.19 |
| T598I | 1.52 | 0.19 |
| A727V | 1.52 | 0.19 |
| K600E (A1798G) | 1.52 | 0.19 |

| | | |
|---|---|---|
| L596R (Exon 15) | 1.52 | 0.19 |
| V599D | 1.51 | 0.19 |
| V599K | 1.51 | 0.19 |
| V599R | 1.51 | 0.19 |
| TP53 | 1.40 | 0.34 |
| 4-Exon (c.102C>A) | 1.40 | 0.34 |
| 4-Exon (c.108G>A) | 1.40 | 0.34 |
| 4-Exon (P47S) | 1.41 | 0.32 |
| 4-Exon (R72P) | 1.41 | 0.32 |
| 6-Exon (c.639A>G) | 1.40 | 0.32 |
| 6-Exon (V217M) | 1.40 | 0.33 |
| 10-Exon (G630A) | 1.41 | 0.34 |

**Table S2(b).** Computed values of fractal dimension and lacunarity coefficient corresponding to the analyzed DNA walks of respective wild-type and mutated oncogenes and tumor-suppressor genes displayed in Fig. 6

| Gene | $\alpha_{max}$ | $\alpha_{min}$ | $\Delta\alpha$ |
|---|---|---|---|
| CRTC1 | 3.07 | 2.00 | 1.07 |
| MAML2 | 2.99 | 2.00 | 0.99 |
| CRTC1-MAML2 | 3.01 | 2.00 | 1.01 |
| SS18 | 2.95 | 1.99 | 0.96 |
| SSX1 | 3.35 | 1.99 | 1.36 |
| SS18-SSX1 | 3.03 | 1.99 | 1.04 |
| SSX2 | 3.41 | 1.99 | 1.42 |
| SS18-SSX2 | 3.32 | 1.99 | 1.33 |
| KRAS | 3.18 | 1.99 | 1.19 |
| G12C | 3.25 | 1.99 | 1.26 |
| G12D | 3.28 | 1.99 | 1.29 |
| G12V | 3.28 | 1.99 | 1.29 |
| PIK3CA | 3.23 | 1.99 | 1.24 |
| Q546K (BEAMing Assay Only) | 3.23 | 1.99 | 1.24 |
| E545K (Exon 9) | 3.23 | 1.99 | 1.24 |
| E545G (Exon 9) | 3.24 | 1.99 | 1.25 |
| E542K (Exon 9) | 3.23 | 1.99 | 1.24 |
| C420R | 3.23 | 1.99 | 1.24 |
| E545A (Tissue Assay Only) | 3.23 | 1.99 | 1.24 |
| H1047R (Exon 20) | 3.23 | 1.99 | 1.24 |
| H1047L (Exon 20) | 3.24 | 1.99 | 1.25 |
| SFPQ | 3.17 | 2.00 | 1.17 |
| TFE3 | 3.33 | 1.99 | 1.34 |
| SFPQ-TFE3(1) | 3.13 | 2.00 | 1.13 |
| SFPQ-TFE3(2) | 3.17 | 2.00 | 1.17 |
| TMPRSS2 | 3.38 | 1.98 | 1.40 |
| ETV4 | 3.46 | 1.99 | 1.47 |
| TMPRSS2-ETV4 | 3.57 | 1.97 | 1.59 |
| PTPRK | 3.29 | 1.99 | 1.31 |
| RSPO3 | 3.25 | 1.99 | 1.25 |
| PTPRK-RSPO3(1) | 3.09 | 1.99 | 1.10 |
| PTPRK-RSPO3(2) | 3.09 | 1.99 | 1.10 |
| MYB | 3.19 | 2.00 | 1.19 |
| NFIB | 3.31 | 1.99 | 1.32 |
| MYB-NFIB | 3.06 | 2.00 | 1.06 |
| PML | 3.32 | 1.98 | 1.33 |
| RARA | 3.33 | 1.99 | 1.34 |
| PML-RARA(1) | 3.36 | 1.99 | 1.37 |
| PML-RARA(2) | 3.36 | 1.99 | 1.37 |
| PML-RARA(3) | 3.36 | 1.99 | 1.37 |
| PML-RARA(4) | 3.36 | 1.99 | 1.37 |
| EWSR1 | 3.23 | 2.00 | 1.23 |
| ATF1 | 3.35 | 1.99 | 1.36 |
| EWSR1-ATF1(1) | 3.30 | 2.00 | 1.30 |
| EWSR1-ATF1(2) | 3.30 | 2.00 | 1.30 |
| EWSR1-ATF1(3) | 3.14 | 2.00 | 1.14 |
| FLI1 | 3.21 | 1.99 | 1.22 |
| EWSR1-FLI1(1) | 3.19 | 2.00 | 1.19 |
| EWSR1-FLI1(2) | 3.12 | 2.00 | 1.12 |
| EWSR1-FLI1(3) | 3.16 | 2.00 | 1.16 |
| EWSR1-FLI1(4) | 3.21 | 2.00 | 1.21 |
| ALK | 3.33 | 1.99 | 1.34 |
| EML4 | 3.14 | 1.99 | 1.15 |
| EML4-ALK(1) | 3.24 | 1.99 | 1.25 |

| | | | |
|---|---|---|---|
| EML4-ALK(2) | 3.21 | 1.99 | 1.22 |
| EML4-ALK(3) | 3.22 | 1.99 | 1.23 |
| BAG4 | 3.37 | 1.99 | 1.38 |
| FGFR1 | 3.38 | 1.98 | 1.40 |
| BAG4-FGFR1 | 3.28 | 1.98 | 1.29 |

**Table S3(a).** Computed minimum and maximum values of Hölder exponents and the multifractality strength corresponding to the analyzed DNA walks of respective wild-type and mutated oncogenes illustrated in Fig. 9 and 10, respectively

| Gene | $\alpha_{max}$ | $\alpha_{min}$ | $\Delta\alpha$ |
|---|---|---|---|
| KIF5B | 3.11 | 2.00 | 1.11 |
| MET | 3.36 | 1.98 | 1.39 |
| KIF5B-MET | 3.10 | 2.00 | 1.10 |
| c.3007 | 3.36 | 1.98 | 1.39 |
| c.2888-5_2944 DEL | 3.32 | 1.97 | 1.35 |
| c.3001_3021 DEL | 3.33 | 1.98 | 1.35 |
| Exon 14 DEL | 3.33 | 1.97 | 1.36 |
| EGFR | 3.17 | 1.99 | 1.18 |
| RAD51 | 3.35 | 1.99 | 1.36 |
| EGFR-RAD51 | 3.22 | 1.98 | 1.24 |
| T790M | 3.19 | 1.99 | 1.20 |
| L858R | 3.17 | 1.99 | 1.18 |
| G719A | 3.19 | 1.99 | 1.20 |
| S768I | 3.20 | 1.99 | 1.11 |
| V441F (VUS) | 3.20 | 1.99 | 1.11 |
| P373L (VUS) | 3.19 | 1.99 | 1.20 |
| L858L (c.2572C>T) | 3.23 | 1.99 | 1.24 |
| L858L (c.2574G>T) | 3.19 | 1.99 | 1.20 |
| L858L (c.2574G>A) | 3.19 | 1.99 | 1.20 |
| L858L (c.2574G>C) | 3.17 | 1.99 | 1.18 |
| Exon 19 DEL | 3.24 | 1.99 | 1.25 |
| Exon 20 INS | 3.18 | 1.99 | 1.19 |
| ESR1 | 3.28 | 1.99 | 1.30 |
| E380Q | 3.28 | 1.99 | 1.30 |
| S463P | 3.25 | 1.99 | 1.26 |
| V534E | 3.28 | 1.99 | 1.30 |
| P535H | 3.26 | 1.99 | 1.27 |
| L536P | 3.28 | 1.99 | 1.30 |
| L536R | 3.30 | 1.99 | 1.31 |
| L536H | 3.26 | 1.99 | 1.27 |
| L536Q | 3.26 | 1.29 | 1.27 |
| Y537S | 3.26 | 1.99 | 1.27 |
| Y537N | 3.36 | 1.99 | 1.37 |
| Y537C | 3.37 | 1.99 | 1.38 |
| D538G | 3.26 | 1.99 | 1.27 |
| TP53 | 3.29 | 1.99 | 1.30 |
| 4-Exon (c.102C>A) | 3.29 | 1.99 | 1.30 |
| 4-Exon (c.108G>A) | 3.30 | 1.99 | 1.31 |
| 4-Exon (P47S) | 3.30 | 1.99 | 1.31 |
| 4-Exon (R72P) | 3.31 | 1.99 | 1.32 |
| 6-Exon (c.639A>G) | 3.30 | 1.99 | 1.31 |
| 6-Exon (V217M) | 3.29 | 1.99 | 1.30 |
| 10-Exon (G630A) | 3.25 | 1.99 | 1.26 |
| KIAA1549 | 3.20 | 1.99 | 1.21 |
| BRAF | 3.20 | 1.99 | 1.21 |
| KIAA1549-BRAF | 3.05 | 2.00 | 1.05 |
| V600E | 3.28 | 1.99 | 1.29 |
| R461I | 3.18 | 1.99 | 1.19 |
| R462I | 3.20 | 1.99 | 1.21 |
| I462S | 3.20 | 1.99 | 1.21 |
| I463S | 3.18 | 1.99 | 1.19 |
| G463E | 3.20 | 1.99 | 1.21 |
| G464E | 3.21 | 1.99 | 1.22 |
| G463V | 3.20 | 1.99 | 1.21 |

| | | | |
|---|---|---|---|
| G464V | 3.20 | 1.99 | 1.21 |
| G465A/G466A | 3.20 | 1.99 | 1.21 |
| G465E/G466E | 3.20 | 1.99 | 1.21 |
| G465V (Exon 11) | 3.18 | 1.99 | 1.19 |
| G468A+1 | 3.20 | 1.99 | 1.21 |
| G468E | 3.18 | 1.99 | 1.19 |
| N580S | 3.20 | 1.99 | 1.21 |
| E585K | 3.21 | 1.99 | 1.22 |
| D593V | 3.18 | 1.99 | 1.19 |
| F594L | 3.18 | 1.99 | 1.19 |
| G595R | 3.20 | 1.99 | 1.21 |
| L596V | 3.18 | 1.99 | 1.19 |
| T598I | 3.18 | 1.99 | 1.19 |
| A727V | 3.22 | 1.99 | 1.23 |
| K600E (A1798G) | 3.20 | 1.99 | 1.21 |
| L596R (Exon 15) | 3.18 | 1.99 | 1.19 |
| V599D | 3.21 | 1.99 | 1.22 |
| V599K | 3.19 | 1.99 | 1.20 |
| V599R | 3.18 | 1.99 | 1.19 |

**Table S3(b).** Computed minimum and maximum values of Hölder exponents and the multifractality strength corresponding to the analyzed DNA walks of respective wild-type and mutated oncogenes and tumor-suppressor genes displayed in Fig. 11

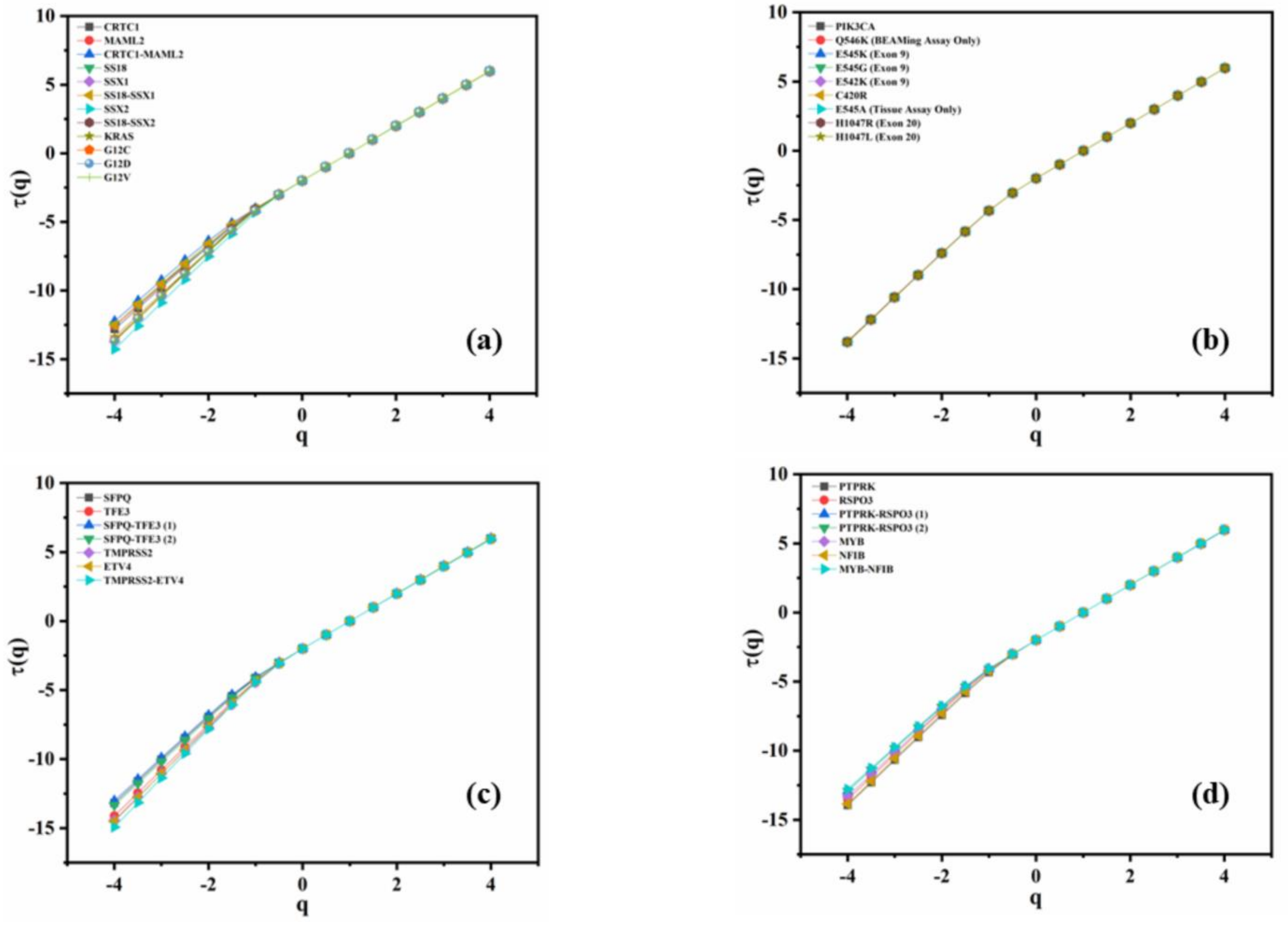


**Fig. S1** Linear behavior of mass exponent with moment for **(a)** wild-type CRTC1, MAML2, SS18, SSX1, SSX2, and KRAS along with their respective fusion and point mutations, **(b)** wild-type PIK3CA and its point-mutated forms, **(c)** wild-type SFPQ, TFE3, TMPRSS2, and ETV4 with their respective fusion variant, and **(d)** wild-type PTPRK, RSPO3, MYB, and NFIB with their particular fused forms

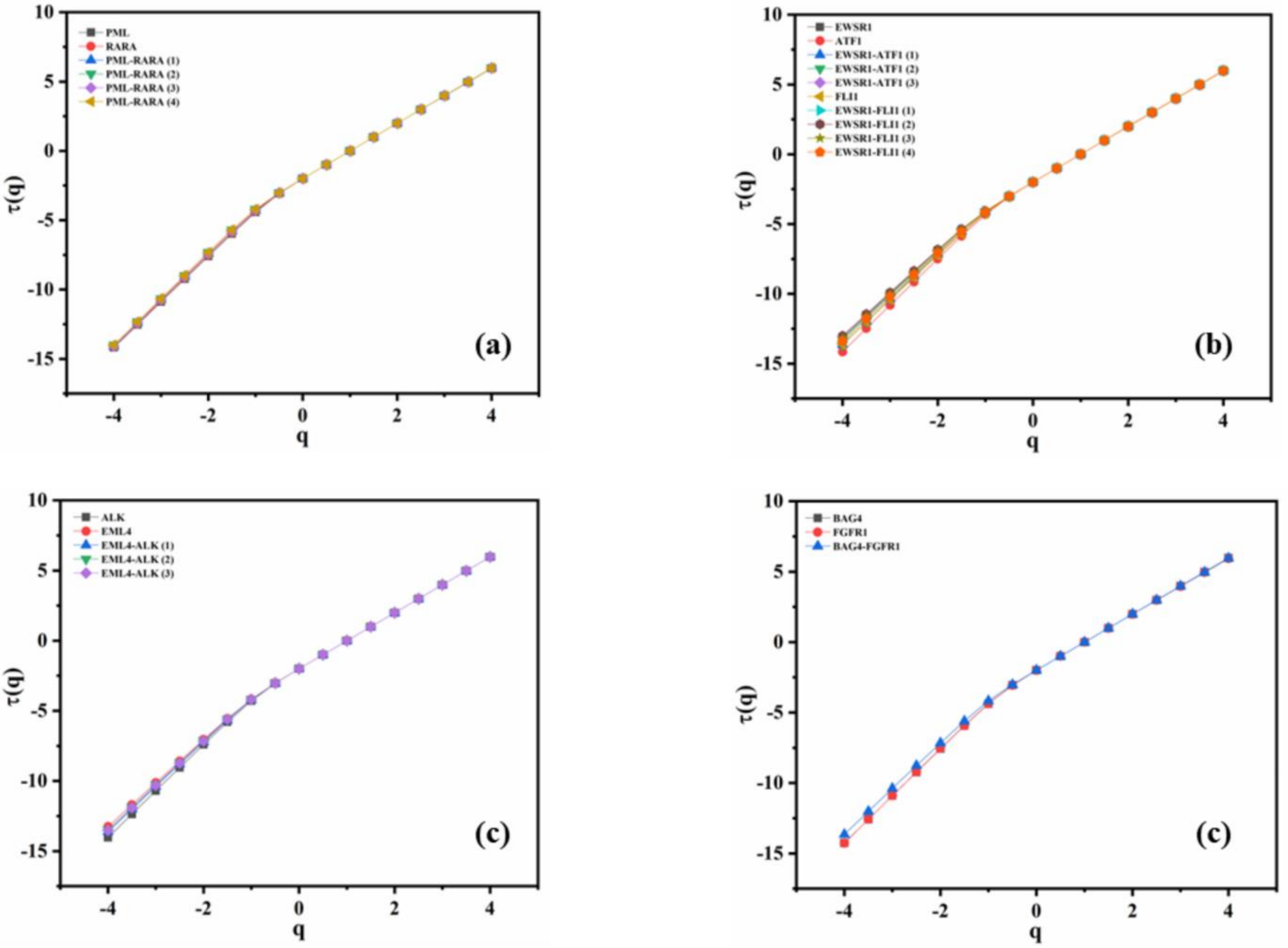


**Fig. S2** Linear behavior of mass exponent with moment for **(a)** wild-type PML and RARA along with their respective fusion form, **(b)** wild-type EWSR1, ATF1, and FLI1 along with individual fusion mutation, **(c)** wild-type ALK and EML4 along with their respective fusion form, and **(d)** wild-type BAG4 and FGFR1 and their fusion form

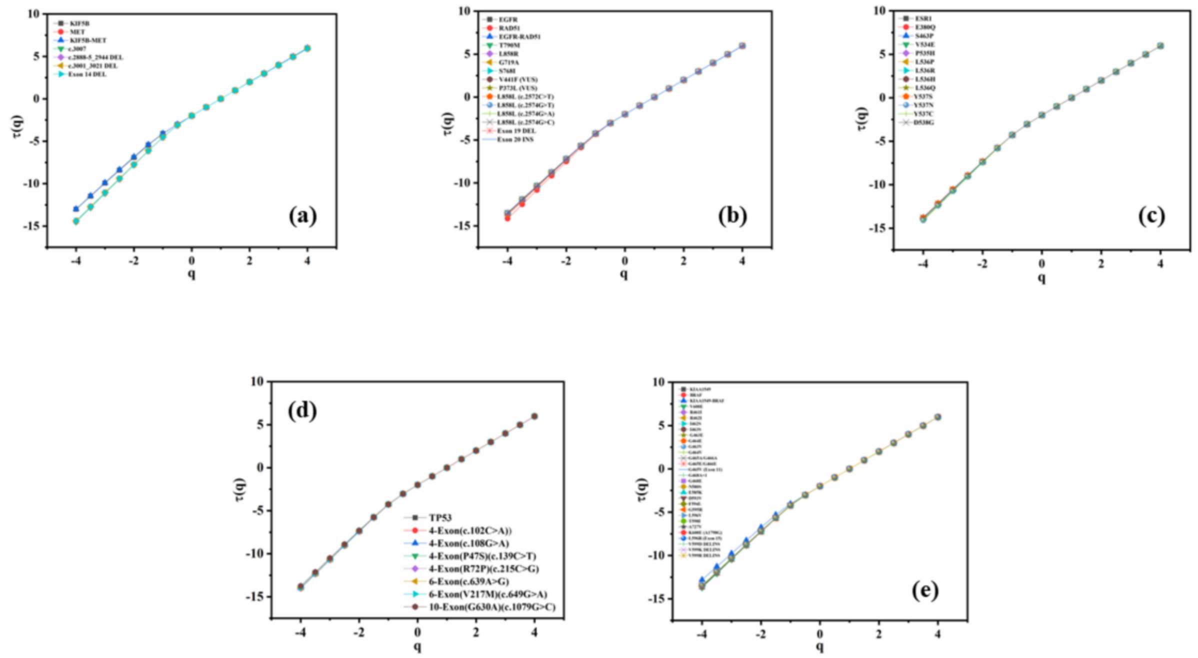


**Fig. S3** Linear scaling of mass exponent with moment for **(a)** wild-type KIF5B and MET along with their respective fusion, and point/deletion mutation in MET **(b)** wild-type EGFR and RAD51 along with individual fusion mutation and point mutations in EGFR **(c)** wild-type ESR1 along with its point mutations, **(d)** wild-type TP53 and its polymorphic forms, and **(e)** KIAA1549 and BRAF along with their individual fusion mutation and point/DELINS mutations in BRAF
